# Expanded Multiplexing on Sensor-Constrained Microfluidic Partitioning Systems

**DOI:** 10.1101/2022.12.23.521805

**Authors:** Pavan K. Kota, Hoang-Anh Vu, Daniel LeJeune, Margaret Han, Saamiya Syed, Richard G. Baraniuk, Rebekah A. Drezek

## Abstract

Microfluidics can split samples into thousands or millions of partitions such as droplets or nanowells. Partitions capture analytes according to a Poisson distribution, and in diagnostics, the analyte concentration is commonly calculated with a closed-form solution via maximum likelihood estimation (MLE). Here, we present a generalization of MLE with microfluidics, an extension of our previously developed Sparse Poisson Recovery (SPoRe) algorithm, and an *in vitro* demonstration with droplet digital PCR (ddPCR) of the new capabilities that SPoRe enables. Many applications such as infection diagnostics require sensitive detection and broad-range multiplexing. Digital PCR coupled with conventional target-specific sensors yields the former but is constrained in multiplexing by the number of available measurement channels (e.g., fluorescence). In our demonstration, we circumvent these limitations by broadly amplifying bacteria with 16S ddPCR and assigning barcodes to nine pathogen genera using only five nonspecific probes. Moreover, we measure only two probes at a time in multiple groups of droplets given our two-channel ddPCR system. Although individual droplets are ambiguous in their bacterial content, our results show that the concentrations of bacteria in the sample can be uniquely recovered given the pooled distribution of partition measurements from all groups. We ultimately achieve stable quantification down to approximately 200 total copies of the 16S gene per sample, enabling a suite of clinical applications given a robust upstream microbial DNA extraction procedure. We develop new theory that generalizes the application of this framework to a broad class of realistic sensors and applications, and we prove scaling rules for system design to achieve further expanded multiplexing. This flexibility means that the core principles and capabilities demonstrated here can generalize to most biosensing applications with microfluidic partitioning.

## 1 Introduction

The advent of microfluidics in biosensing has led to portable, cost-effective, and automated assays to be performed on simple chips manufactured with the same platforms that spurred the computing revolution [1,2]. However, the core methods of biosensing have largely rested on the paradigm of designing sensors that are each specific to a target analyte. For situations in which many target analytes must be considered, this one-to-one principle scales poorly: many sensors must be embedded on a single device, samples must be concentrated enough such that a representative subsample can be applied to each sensor, and cross-reactivity of sensors and analytes scales combinatorially [3, 4]. Our motivating application is in bacterial and fungal infection diagnostics where one or a handful out of hundreds of plausible pathogens may be responsible for a patient’s condition, but samples may exhibit very low microbial concentrations [5, 6]. For instance, a milliliter of blood can have as low as one colony forming unit (viable cell) or on the order of 10^2^ to 10^3^ equivalent genomic copies of microbial DNA [7].

Scalable coverage of many analytes requires circumventing the one-to-one scaling of specific sensors. *Nonspecific* sensing modalities that each generate measurements from multiple analytes are necessary for higher order multiplexing provided a post-processing method for inferring the presence or quantities of individual analytes. For nucleic acid diagnostics, DNA sequencing is often the method-of-choice. Metagenomic sequencing analyzes the contents of virtually with raw sequence reads analyzed and interpreted with bioinformatics, but this approach has limited sensitivity in the presence of high background such as host DNA in blood [8]. Amplicon sequencing is an alternative for microbial diagnostics in both microbiome analysis and infections [9,10]. These approaches conduct PCR on ribosomal RNA genes (e.g., 16S for bacteria, 18S or 28S for fungal, among others), that are flanked by conserved regions for priming and exhibit internal sequence differences for taxonomic discrimination [9, 11]. While sequencing has made strides in clinical practice, its expense, required expertise, and complex workflows have hindered its routine use [12, 13].

Another category of nonspecific sensing involves “fingerprinting” where a general sensing modality assigns unique signatures to target analytes such that an unknown sample can be read and matched against a database of analytes. Spectroscopic methods are common in this class, profiling wide ranges of proteins, metabolites, and cells. In clinical infections, mass spectrometry has been applied to rRNA amplicons [14] although its use to identify clinical isolates from positive culture is gaining much more traction [15, 16]. However, a major limitation of these approaches is the difficulty in analyzing mixtures of analytes [17, 18]. For mass spectrometry, this limitation manifests as a need to analyze clinical isolates from polymicrobial samples one at a time.

Microfluidic partitioning technologies offer an avenue for the high throughput, sensitive, and quantitative characterization of heterogenous samples via a fingerprinting approach [17]. These systems split an initial sample into thousands or millions of partitions such as droplets or nanowells [19]. If the analytes are at a limiting dilution, most partitions will be empty with an occasional analyte isolated in its own partition. Formally, analytes are captured according to a Poisson distribution and the Poisson rate parameter *λ*_*n*_ for analyte *n* dictates the probability of capturing nonnegative integer copies (*x*_*n*_) of the analyte in a given partition with 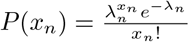 [20,21]. Among *N* total analytes indexed by *n* that are distributed independently among the partitions, single-analyte capture is very likely if ∑_*n*_ *λ*_*n*_ < 0.1 with most partitions remaining empty. This “digital microfluidics” approach guides much of the research in single-cell and single-molecule analysis.

Given the probabilistic isolation of individual analytes, nonempty partitions can be classified one at a time against a database. Researchers have demonstrated classification with high resolution melt (HRM) curve analysis of individual 16S gene amplicons captured in droplets [22]. Also, surface enhanced Raman spectroscopy (SERS) of isolated bacterial cells [23] can assign unique spectra to species, and digital SERS with microfluidic capture has been proposed [17]. However, all microfluidic partitioning technologies rest on the assumption that droplets must be individually classified. From a data perspective, these systems are dependent on reliable decision boundaries between *N* analyte classes which makes them highly sensitive to measurement noise [24]. Acquiring enough information from each partition for reliable classification limits the throughput of acquiring partition measurements, and therefore, the volume of sample that can be analyzed [25]. Moreover, in diagnostics, sample concentrations are rarely known *a priori*. Samples with higher concentrations can result in multi-analyte capture in the same partition, causing errors in classification approaches that assume single-analyte capture.

Our group recently coupled advances in microfluidic partitioning with ideas from compressed sensing to address these challenges. Compressed sensing (CS) seeks to infer *sparse* signals efficiently: faster or with fewer sensors [26, 27]. In biosensing, samples are sparse when among many possible analytes, only a handful are present in any given sample. For instance, a patient could be infected with any of hundreds of pathogens, but only one or a few are responsible for the current infection [28]. In this application, CS is analogous to quantifying analyte fingerprints from mixed measurements [29]. Our group recently developed new theory and a new Sparse Poisson Recovery (SPoRe) algorithm that couples principles of CS with microfluidic partitioning [30]. SPoRe performs maximum likelihood estimation (MLE) via gradient ascent over a generalized likelihood function. While we chose to explore this direction because microfluidics approaches the limit of sensitivity through single-molecule analysis, we also found fundamental advantages from a signal processing perspective. Most notably, leveraging the Poisson-distributed capture of analytes enables improved rates of multiplexing (fewer sensors, more analytes), tolerates multi-analyte capture in the same partition, withstands very high measurement noise, and can enable partial fingerprints to be captured separately in sensor-constrained systems.

This latter concept of *asynchronous fingerprinting* enables high-throughput, sensor-constrained microfluidics systems to achieve both sensitive detection and efficient multiplexing of analytes. The key insight is that individual partitions can be entirely ambiguous in their analyte content, but the *distribution* of all partition measurements can be used to solve for the analyte concentrations. In this work, we first extend our statistical theory to cover a broad class of realistic sensors Next, we present the first *in vitro* demonstration of our framework towards bacterial infection diagnostics, quantifying twelve bacterial species at the genus level with only five orthogonal DNA probes in two-channel droplet digital PCR (ddPCR). We selected species based on high prevalence in bacterial diagnostics and cause for concern due to growing drug resistance [31–33]. We characterize the performance of our assay with 18 samples each with a mixture of 2-4 bacteria, demonstrating accurate polymicrobial quantification down to approximately 200 total copies of the 16S gene. Finally, we show how our probabilistic framework enables the flagging of samples that contain 16S barcodes outside of the designed panel. Our goal is that the promising practical results of our demonstration motivate further theoretical research, a refinement of our particular assay towards scalable infection diagnostics, and broader applications of our new framework to multiplexed biosensing.

## 2 Results

### 2.1 Overview of ddPCR Approach

In ddPCR, microfluidics splits a sample into thousands of droplets to stochastically capture nucleic acids (Fig. 1). Endpoint PCR measurements form binary clusters that indicate the presence or absence of target sequences [34, 35]. We used the Bio-Rad Qx 200 (Bio-Rad Laboratories, Hercules, CA, U.S.A.) which has two fluorescence channels (FAM and HEX) for multiplexed PCR with hydrolysis probes.

**Figure 1:**
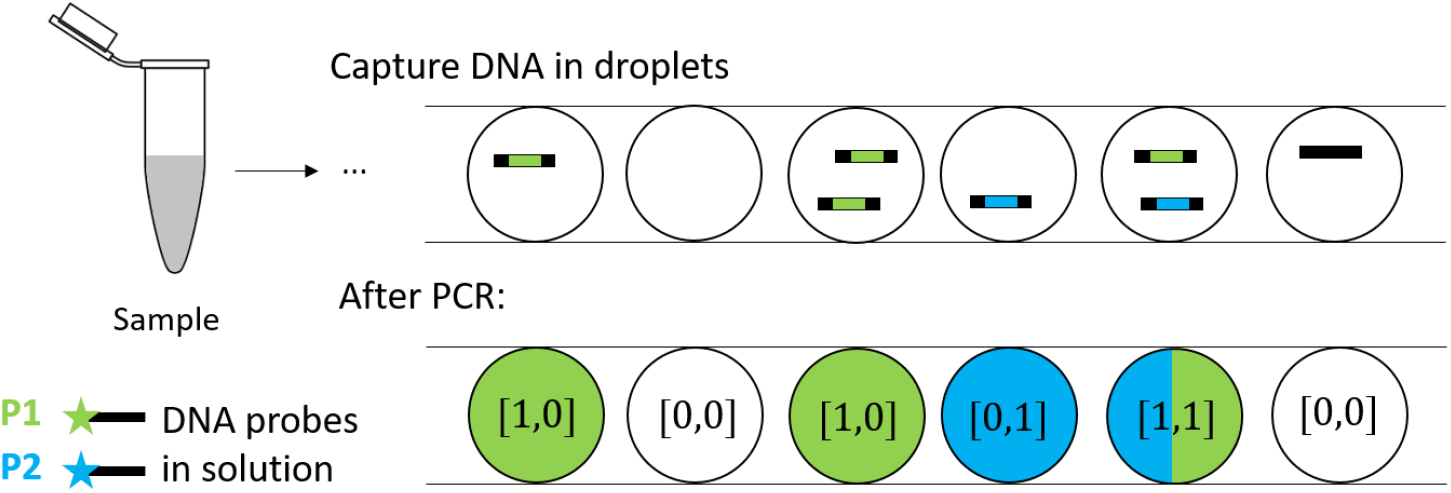
Summary of ddPCR. A sample is split into many droplets, collected, deposited into a ddPCR plate for amplification (not depicted), and read after PCR. Endpoint fluorescence measurements indicate the presence or absence of target nucleic acids in droplets. Black rectangles indicate PCR amplicons, and green or blue rectangles correspond with complementary sequences of the HEX and FAM probes, respectively, in the amplicons. Droplets with amplicons without a probe binding site elicit minimal fluorescence, ideally clustering with genuinely empty droplets.

We designed nonspecific 11-nucleotide hydrolysis probes spiked with locked nucleic acids (LNAs) that bind to the 16S genes of a panel of 12 bacterial species, detailed in Section 4.2. Briefly, we filter the set of 11-mers to avoid heterodimers and weak mismatches while ensuring a sufficiently high melting temperature. Each 16S gene elicits a binary barcode response to the set of five candidate probes based on the presence or absence of the probe sequences in the gene (Fig. 3a). We used coordinate ascent optimization to select a final probe set that separated the bacteria by genus. Particularly, we group three species of *Staphylococcus* together (*S. aureus, S. epidermidis*, and *S. saprophyticus*) and two species of *Streptococcus* (*S. agalactiae* and *S. pneumoniae*).

There are multiple 16S gene copies in each bacterial genome with slight sequence variability. Although we attempted to design probes such that each genus had a unique, consistent barcode for all copies, *E. cloacae* appeared to exhibit a small proportion of variant barcodes, a fraction which we computed experimentally (Section 4.8). Such variation is likely inevitable especially in larger scale systems but can be accounted for.

We store each pathogen’s fractional barcode distribution across its copies in a column of a matrix **C** (Fig. 2). Note the ordering of barcodes is arbitrary in constructing **C** and that because of only slight variation between 16S copies within a genome [11], **C** is nearly the identity matrix in practice. In the rows of **C**, we also ignore the barcodes that are not elicited by the combination of probes and the bacterial panel. Our optimization estimates the 9-dimensional Poisson parameter vector **λ** that represents 9 analyte concentrations. With 9 unique barcodes and bacterial genera, the term “analytes” could refer to either. If the analytes are the barcodes, then 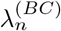 is the concentration of the total 16S genes from *any* source bacteria that exhibits the *n*th barcode. If the analytes are the bacteria, then 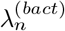 is the concentration of the *n*th bacteria’s 16S genes, regardless of the particular barcodes of individual genes. Because **λ**^(*BC*)^ = **C*λ***^(*bact*)^, our results currently depend on a **C** matrix of rank *N* to readily convert between the barcode concentrations **λ**^(*BC*)^ and the bacterial concentrations **λ**^(*bact*)^. For context-dependent reasons, we make use of both definitions of “analyte”, carefully clarify which we are using at any time, and often drop the superscript notation.

**Figure 2:**
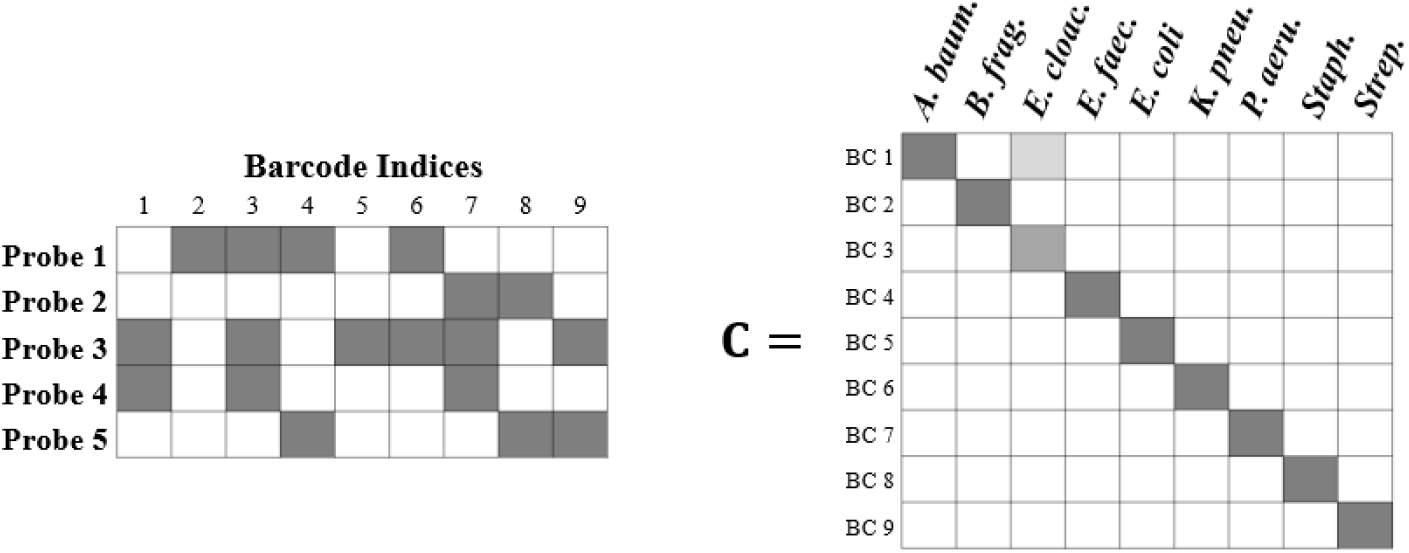
Accounting for barcode variability across copies of the 16S gene. The white values in **C** are zero, with the darkest gray representing 1. Each column contains the proportions of the barcodes that each bacterial taxa’s 16S gene copies exhibit.

**Figure 3:**
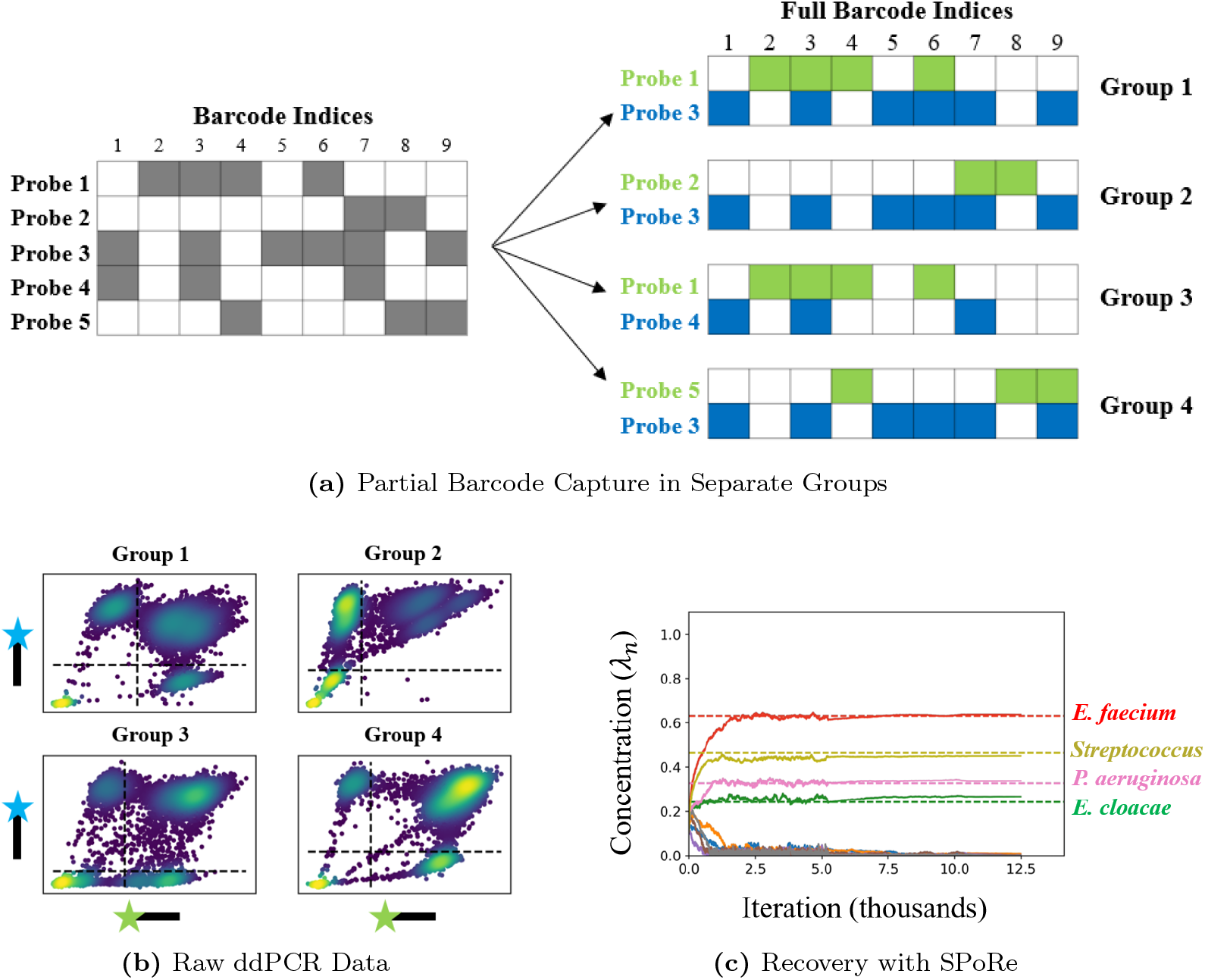
End-to-end example of assay. **(a)** Nonspecific hydrolysis probes react with the 16S genes of multiple bacteria. Probes assign binary barcodes to 16S genes (non-white colors indicates the probe binds to the bacterial gene). All bacteria except *E. cloacae* elicited the same barcode across all gene copies, and *E. cloacae* had an estimated one variant copy (Section 4.8). Due to multiplexing and channel limitations of ddPCR, we capture subsets of the barcodes in four groups of droplets. **(b)** Example of raw data from four groups of droplets, each from the same mixed bacterial sample. Raw data is binarized by manual thresholding, overriding most of the effects due to partition specific competition. **(c)** Sparse Poisson Recovery (SPoRe) algorithm optimizes over all groups simultaneously, accurately reflecting the estimated ground truth (dashed lines).

Ideally, we could estimate **λ**^(*BC*)^ by simply capturing individual 16S genes in droplets with all five probes (Section 5.1). However, the unique clusters only hold for single-gene capture; if multiple genes appear (with distinct sequences) in the same droplet, an effect known as *partition specific competition* (PSC) occurs and fluorescent intensities can decrease [35]. Second, unique clusters for every barcode cannot scale beyond this study if the eventual goal is to quantify dozens to hundreds of microbes.

Instead, we generate four *sensor groups* of droplets each with a different subset of two probes (Fig. 3a). We call this concept *asynchronous fingerprinting* and describe the allocation of probes to each group in our mathematical theory (Theorem 4.7 in Section 4.1.3). Despite competition effects, raw droplets can still be reasonably thresholded above zero in each channel [36]. Although the 16S barcodes in each droplet is made entirely ambiguous, we infer bacterial concentrations in the sample from the pooled, binarized data from these four groups of droplets. SPoRe essentially finds the solution that best explains the distribution of droplet measurements across the four groups.

### 2.2 Generalized MLE with Microfluidic Partitioning

The standard for quantification in digital microfluidics data is based on MLE [37]. We generalize MLE in our new framework and apply it to ddPCR. Respectively, the general terms used in this section *analyte, partition*, and *measurement vector* correspond with the physical concepts of a barcode or bacterium, a droplet, and the two pre-binarized measurements acquired from each droplet. Section 4.1.1 contains detailed clarification of our mathematical notation.

Let *N* and *D* define the number of unique analytes in the assay and the number of partitions, respectively. We let **x**_*d*_ be an *N* -dimensional nonnegative integer vector representing the quantities of each analyte in partition *d*. We say **λ** is *k*-sparse if *k* elements are nonzero. With microfluidic partitioning, **x**_*d*_ is distributed as Poisson(**λ**) where **λ** is the *N* -dimensional parameter vector that characterizes the rate of capture of each of *N* analytes. Let **y**_*d*_ represent the measurement vector acquired from partition *d* (e.g., in our assay, **y**_*d*_ ∈ {0, 1}^2^). Note that while **y**_*d*_ is observed directly, **λ** must be inferred and **x**_*d*_ is latent. We use an asterisk (**λ**^*^, 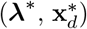) to denote true values and a hat 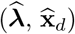 to denote estimates. The goal in MLE is to find an estimate 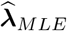 that maximizes the likelihood of the observed measurements:

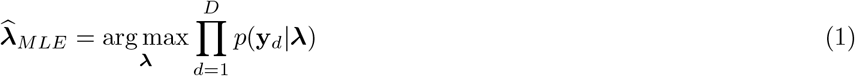

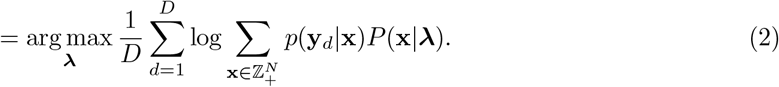

Denoting the likelihood function from the right-hand side of (2) as *ℓ*, the gradient is

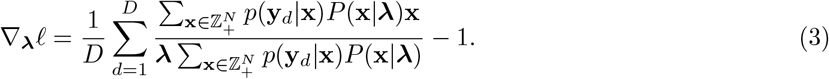

Although fairly obtuse, this equation leads to two commonly observed equations in specialized implementations of MLE that use digital fingerprinting or orthogonal assays (Section 5.3). In our ddPCR assay, droplets may contain multiple gene copies, and no droplet can be assumed to have or lack a particular 16S barcode. Despite this ambiguity, our SPoRe algorithm uses gradient descent to iteratively solve Equation (2).

#### 2.2.1 Modifications to SPoRe Algorithm

We summarize two particular changes to the original SPoRe algorithm with implementation details left to Sections 4.7 and 5.2. First, we previously noted the flexibility of SPoRe for any sensing model. This flexibility arises from the fact that *p*(**y**_*d*_|**x**), the sensing model that maps signals to measurements, is the only term that must be defined for a particular application. We use a simple model for our ddPCR assay. For **y**_*d*_ ∈ {0, 1}^2^ with *M* = 2 for the two fluorescence channels, we say *p*(**y**|**x**) = ∏_*m*_ *p*(*y*_*m*_|**x**) with *p*(*y*_*m*_|**x**) = 1 if **x** has at least one copy of a gene that contains the corresponding probe, with *p*(*y*_*m*_ | **x**) = 0 otherwise. In our implementation, we define the analyte as the bacterial content and account for fractional barcode content in the gradient computations (Section 5.2).

Second, note in Equation (3) that the gradients contain an infinite summation over all of *N* -dimensional nonnegative integer vectors 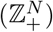. In our earlier work [30], we used Monte Carlo approximations of this gradient on batches of measurements. Here, with the finite measurement space of ddPCR, these gradients can be computed quickly and exactly over all measurements (Section 5.2). We use backtracking line search to speed up convergence of gradient descent to estimate 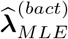. Such accelerations are less feasible if the gradient is loosely approximated. The exact gradient, while cumbersome to derive for the 2-channel ddPCR system, could be similarly calculated for any number of channels and is computationally cheap.

#### 2.2.2 Identifiability of Model

Running gradient descent to convergence will always find some solution, but we need to develop some assurance that it is the correct solution. Proving the *identifiability* of a model ensures that there is a unique global optimum to the likelihood function given infinite data. Formally, identifiability states that if *p*(**y**| **λ**) = *p*(**y** |**λ**′) for all **y** in each sensor group, then **λ** = **λ**′.

We formally define terms and prove sufficiency conditions for identifiability in Section 4.1. Briefly, we find that **λ** can be inferred uniquely if the sensing functions that dictate the mapping of **x** to **y** are monotonic. Monotonic functions do not change direction in output; in biosensing, most outputs *increase* with increases in input and vice versa such that the sensors are monotonic *increasing*. Any additional analyte should not decrease the output measurement. We also assert that if one copy of an analyte does not yield a nonzero measurement, then the analyte is considered *nonresponsive* such that its content in a partition has no influence on the measurement. Lastly, we also impose a system-wide condition called *fingerprint equivalence* which (informally) states that analytes with the same single-molecule fingerprints in a given sensing group behave interchangeably.

To align with these conditions in our proofs, we define the analytes as the barcodes. Under reasonable PSC effects (e.g., not an overwhelming diversity of 16S genes in any given droplet) the addition of new barcodes to a droplet cannot reduce the binarized measurements (monotonicity). The binary data is determined by the presence of a 16S gene with a complementary probe sequence; without such a binding site, the gene is nonresponsive. Lastly, gene copies with the same combination of probe binding sites are interchangeable (fingerprint equivalence). Theorem 4.7 proves the sufficiency of these conditions for **λ**^(*BC*)^ = **λ**^*′*(*BC*)^, but with *rank*(**C**) = *N*, **λ**^(*BC*)^ = **λ**^*′*(*BC*)^ ⇒ **λ**(*bact*) = **λ**^*′*(*bact*)^.

Here, we provide an informal explanation and concrete example of the result of Theorem 4.7 which defines a matrix **Z**^(*g*)^ whose rows indicate the positions of analytes with equal, nonzero fingerprint responses in the sensor group *g*. Stacking these matrices for each *g* yields a matrix **Z**, and if *rank*(**Z**) = *N*, then the system is identifiable. Figure 4 illustrates this process for our particular assay. With two binary measurements per group, there are 2^2^ −1 = 3 nonzero barcode measurements. For instance, note that in Group 1, original barcode indices 2 and 4 share a [1,0] response, yielding the first row of **Z**^(1)^. Each group contributes three rows to the matrix **Z**.

**Figure 4:**
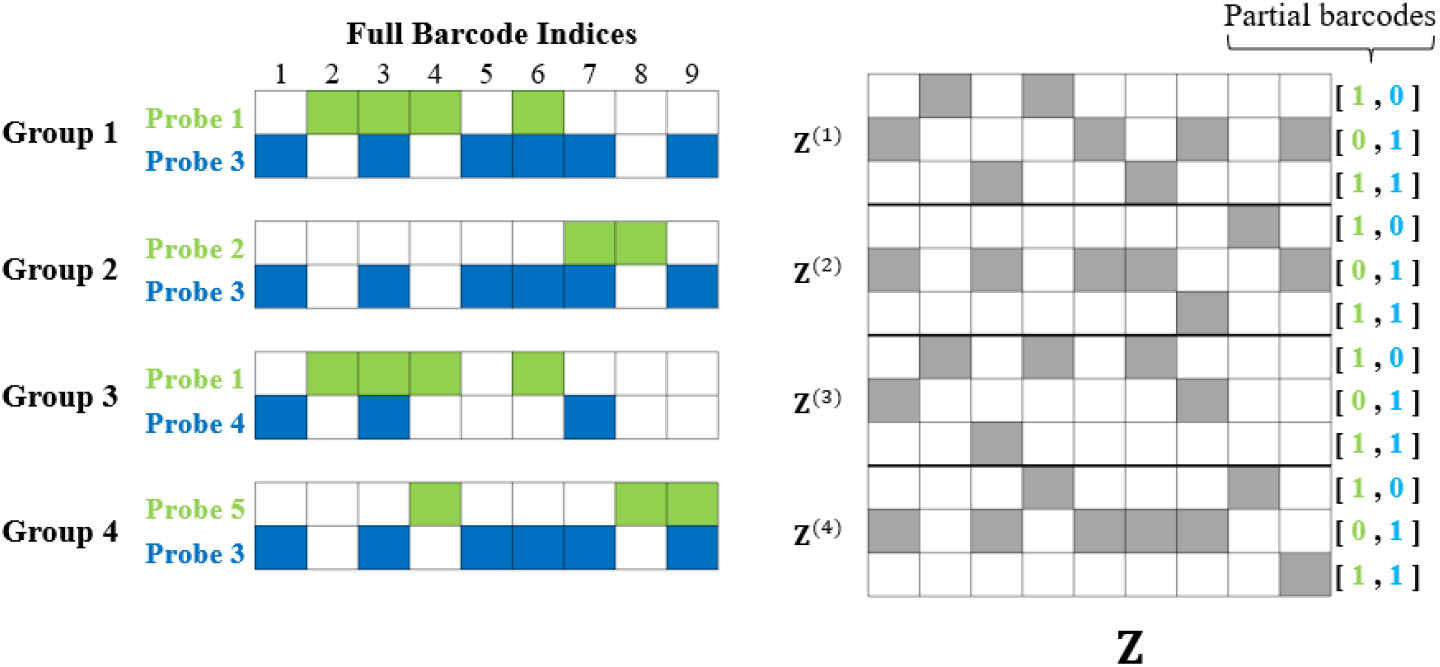
Formation of the linear system matrix **Z** that verifies identifiability for our assay. Non-white squares are 1, and white squares are zero. Each group contributes three rows to **Z** as described in Theorem 4.7 in Methods 4.1.3, and *rank*(**Z**) must be *N*.

This result implies that we cannot arbitrarily assign probes to each group, and interestingly, using probes in multiple groups can be beneficial by adding rows to **Z**. It is not sufficient to simply capture each probe’s information in at least one group. Also, for *rank*(**Z**) to be *N*, we can derive a simple rule of thumb for binary ddPCR with *M* channels: *G*(2^*M*^ − 1) ≥ *N* is necessary for the conditions of Theorem 4.7. Although we had access to two-channel Bio-Rad Qx200, this result also indicates promise for applying our framework to digital PCR systems with more than two channels: up to *N* = 15*G* analytes on the 4-channel QuantStudio Absolute Q (ThermoFisher Scientific, Waltham, MA), or up to 63*G* analytes on new six-channel systems from Bio-Rad and Roche (Basel, Switzerland). Note that we do not intend to rank these instruments, as other factors such as the volume that can be processed, the number of partitions that can be generated, automation of workflows, etc. require application-specific consideration.

However, the implications of identifiability must be approached with caution. From an optimization perspective, identifiability simply implies that given *infinite* data (*D* → ∞) in each group, the true **λ**^*^ is the global optimum of the likelihood function. Our results fall short of a recovery guarantee for a finite *D* partitions. In our earlier work [30], we derived the insight that less sparse **λ**^*^ (more analytes with nonzero quantities) necessitate more partitions for stable recovery. Nonetheless, in contrast to typical applications of CS, there is no explicit maximum for the sparsity level as any **λ** is identifiable under our result.

### 2.3 Demonstration of Polymicrobial Diagnostics

We tested SPoRe’s ability to quantify bacterial loads in mixed samples of purified genomic DNA. We used reference wells with individual bacterial dilutions to estimate the ground truth concentration and assist with manual thresholding (Section 4.6) to binarize the data. We passed this pre-binarized data to SPoRe.

Figure 5a illustrates the quantitative results. For general performance evaluation on polymicrobial samples, we use the cosine similarity metric to capture our framework’s concordance with the true relative abundances of bacteria in the sample. We find an average cosine similarity of 0.96, indicating our ability to very reliably capture the dominant bacteria in a sample while making some errors on the relatively less abundant bacteria. These errors are further characterized in Fig. 5b, which indicates that relative error in the estimated relative abundances decreases for higher relative abundance bacteria. Note that the color of the datapoints in Fig. 5b indicate the absolute abundance of the bacterium, matching the colormap of Fig. 5a. Some bacteria at low absolute abundances but high relative abundances achieve low relative error. Of course, 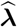 estimates absolute abundance. Sweeping a global threshold on 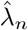 to make a binary call on bacterial presence yields a receiver operating characteristic curve with an AUC of 0.966 (Fig. 5c, and a downward trend in AUC when increasing *k*. From our prior theoretical work, less sparse samples are subject to higher estimation variance which may explain this effect [30]. A fixed threshold of *λ*_*n*_ = 0.15 achieves an overall sensitivity of 94% and specificity of 95%.

**Figure 5:**
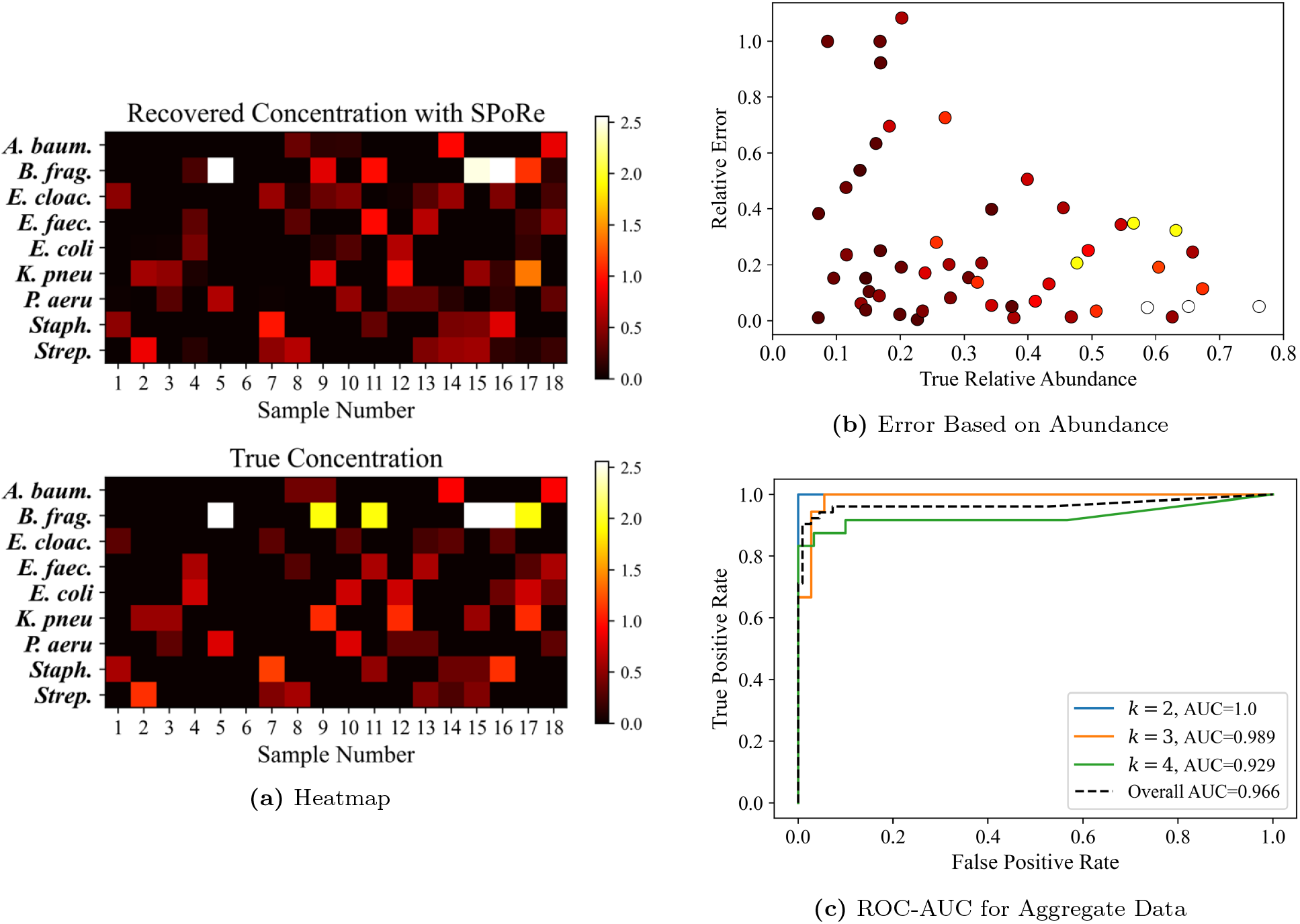
(a) Signal recovery results against the estimated ground truth. All colors are scaled against the maximum estimated ground truth concentration of *λ*^*^ = 2.56 for Concentration 1 of *B. fragilis*. Sample 6 is a negative control with no baterial genomic DNA added. (b) Relative error of the estimated relative abundance versus true relative abundance. Datapoints use the same colormap in (a) to indicate the estimated absolute abundance of the bacterium. (c) Receiver operating characteristic curves and their area under the curves (AUC) on aggregate data within different sparsity levels and across all samples.

We continue with an extensive investigation on the errors in recovery. Solving an optimization problem can risk returning a locally optimal solution. However, we find that the likelihood of the converged solution is higher than that of the supposed ground truth (Fig. 6), so it is unlikely that local optima explain our discrepancies. Differing in concentration estimates for the bacteria truly in the sample is to be expected, as some of this error will arise simply from pipetting volume variablity and sampling variability in capturing 16S gene copies.

**Figure 6:**
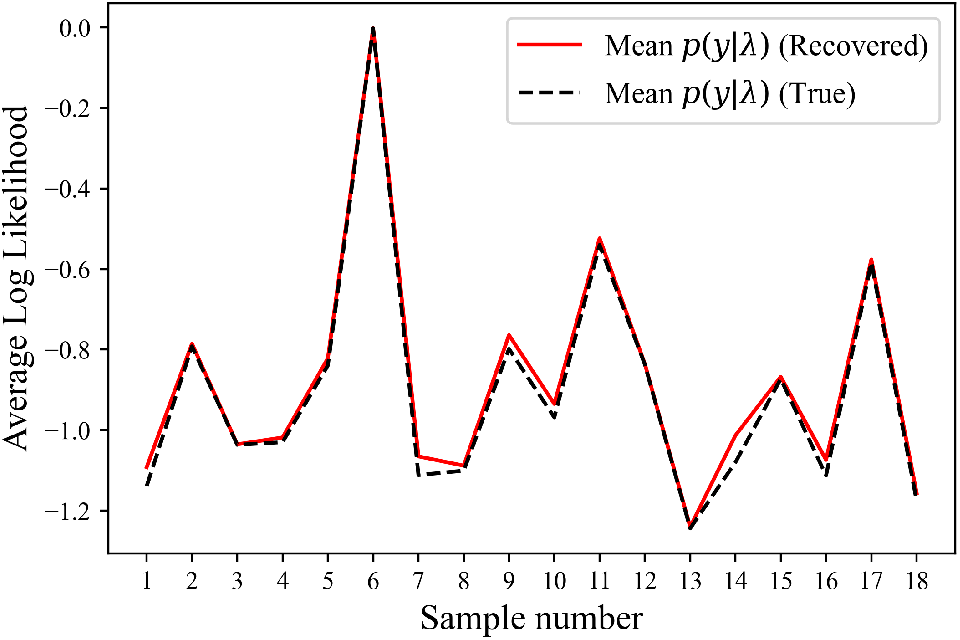
Likelihood comparison of SPoRe’s solution against the estimated ground truth. SPoRe’s solution exhibits higher average likelihood for the pre-binarized data that it is given.

However, in some samples, SPoRe misses a bacteria of low abundance (a false negative) while including a bacteria that is absent in the sample (a false positive). We hypothesize that these errors may be due to imperfect thresholding of our data that could easily be misclassifying some droplets. Because we pass pre-binarized data to SPoRe, mistakes in thresholding propagate to SPoRe; that is, given warped data, SPoRe will return a warped solution which could appear to have higher mean likelihood than the estimated ground truth (6). Some degree of droplet misclassification is very likely with our current assay for three reasons illustrated in Figure 10. First, our measurements currently exhibit very high droplet *rain* which we suspect is due to our attempts to amplify a very long sequence: the 16S gene is roughly 1500 base pairs, while Bio-Rad recommends an amplicon of 75-200 base pairs. Second, PSC may slightly blur the boundaries between the typical binary clusters in droplets with more than one amplicon. Third, the negative droplets also exhibit some “lean” and “lift” most likely due to partial probe interactions with amplicons containing slightly mismatched sequences. Overall, these combined effects squeeze the boundary in which we can draw thresholds. While these effects are common and have some popular tools to help disambiguate droplets [38, 39], we decided to use manual clustering as these tools are generally not designed for the conditions of our assay.

We designed a simulated experiment to directly evaluate the effect of imperfect binarization of the raw measurements. Given the estimated ground truth concentrations and the droplet counts in each group, we simulated underlying droplet gene content (**X** with **x**_*d*_ ∼ *Poisson*(**λ**^*^)) and the resulting binary measurements using our *p*(**y** | **x**) model. On this simulated data, SPoRe returned virtually perfect solutions with a mean cosine similarity of 0.9997 (Fig. 13). This improved performance indicates that the most likely cause of errors with our thresholded data is due to our model’s imperfect ability to represent the data. This discrepancy is natural given that our model asserts hard thresholds for binarization while the real data does not maintain strict decision boundaries. However, our high cosine similarity with the real data illustrates that our signal reconstruction is robust to some model imperfections in recovering the relatively high abundance bacteria in samples.

### 2.4 Characterization of Limit of Quantification

In infection diagnostics, pathogen loads can vary by several orders of magnitude. Tolerating multi-gene capture reduces the risk that high concentration samples flood a system and allows design flexibility for microfluidics systems with fewer partitions (e.g, smaller form factors with nanowells instead of droplets). We designed samples such that total bacterial concentrations 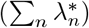 would be between 1 and 5 to illustrate this ability. However, demonstrating this capability on the Bio-Rad Qx200 meant that our samples have 16S concentrations that are unrealistically high for most clinical presentations.

We characterized the limit of quantification in terms of 16S copy counts per sample for partitioning systems that may still result in multi-analyte capture by randomly subsampling our experimental data. For each sample, we subsampled 10%, 1%, 0.1% and 0.01% of the droplet data and passed it to SPoRe. We estimate the 16S copy count in this data as the product of the number of subsampled droplets and the total estimated ground truth concentration 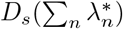. Figure 7 shows how SPoRe maintains strong recovery down to approximately 200 copies of the 16S gene. Depending on the quality of a future upstream microbial DNA isolation procedure, this limit could be potent for many applications in infection diagnostics.

**Figure 7:**
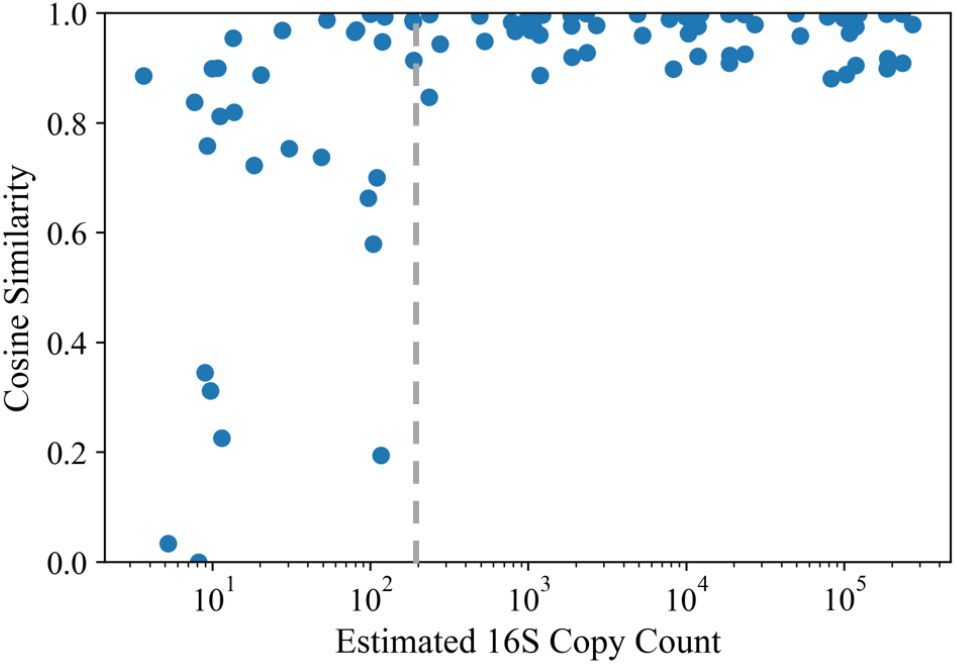
SPoRe’s performance on random subsamples of experimental data. Each sample’s population of pre-binarized droplet measurements was subsampled by a factor of 10^−1^, 10^−2^, 10^−3^, and 10^−4^. SPoRe performed MLE on the subsampled data.

Of course, a final system may have the flexibility to generate many partitions, meaning that at low 16S copy counts, non-empty partitions most likely have only one target molecule captured. While this is uniquely unnecessary with our generalized approach to MLE, low magnitudes of **λ** empirically help recovery [30]. Intuitively, if the same sample is split over more partitions, signal inference can only stand to gain information by capturing measurements from more individual molecules rather than their combined effects. Moreover, in ddPCR, single-molecule capture would avoid PSC altogether such that thresholding may be more reliable.

### 2.5 Flagging of Samples with Unknown Barcodes

Given a set of droplet measurements, MLE will always report some solution even if the sample contains a bacterium with a 16S barcode distribution outside of the panel given to SPoRe. However, this probabilistic approach allows us to assess the quality of the recovered solution and detect such anomalies. Given a recovered 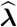, we can characterize the expected distribution of the discrete measurement vectors and perform a *χ*^2^ goodness of fit test between the expected and observed distribution of measurements. A poor match between these distributions would indicate a faulty solution that could be due to an out-of-panel barcode distribution.

We used the *p* value of the *χ*^2^ goodness of fit test as a metric to detect faulty solutions. For each tested polymicrobial sample, we simulated the effect of having an “unknown” bacterium in it by removing each of the correct, present bacteria (one at a time) from SPoRe’s database before running the algorithm. We repeated this process for both the manually thresholded and simulated data. In both cases, the *p* value of the test is a highly reliable metric for flagging samples with out-of-panel bacteria as indicated by the receiver operating characteristic (ROC) curves (Fig. 8). With simulated data, the separation is perfect with an area under the curve (AUC) of 1.0. Indeed, the minimum *p* value for SPoRe on simulated data with the full database of microbes was 0.783, and all cases in which SPoRe was deliberately not given one of the present bacteria in the sample returned *p* = 0. Given a reliable measurement model that corresponds with real-world data, the significant presence of a microbial barcode outside the provided database could be reliably detected with a pass threshold on the *p* value. With manual thresholding, note that small mistakes in binarizing the data may make the observed distribution of measurements improbable for any **λ**. As a result, samples which contain only bacteria in the panel may nonetheless return results that are flagged as faulty. We see this effect in the diminished (but still strong) performance with an AUC of 0.964.

**Figure 8:**
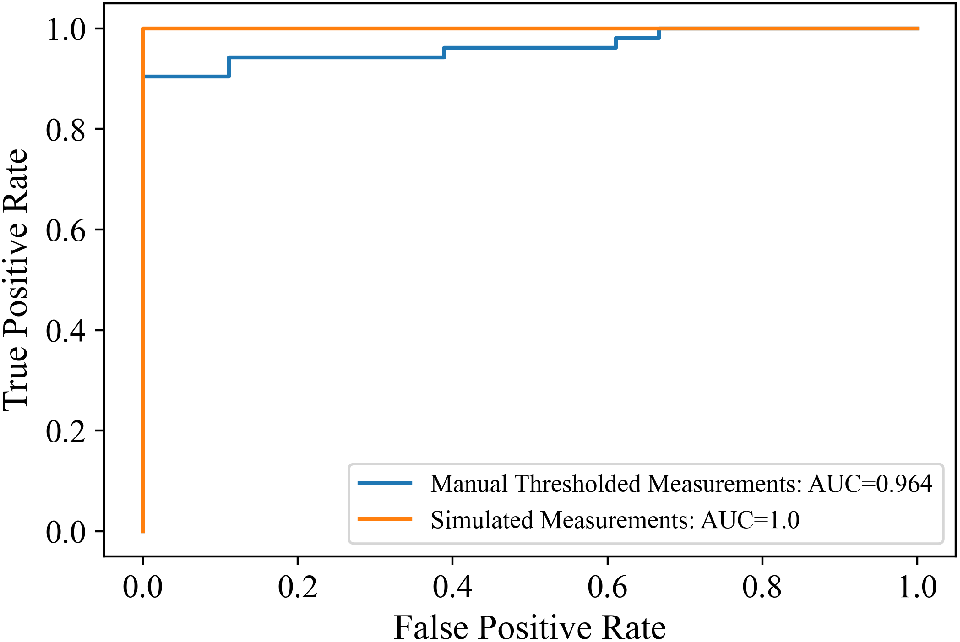
Flagging samples with out-of-panel bacteria. A *χ*-squared goodness of fit test is performed using the distribution of **y** expected given 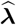 and the observed distribution. The *p* value of the test is used as the metric for determining if a sample has an unknown barcode (e.g., SPoRe was not given one of the present barcodes in the solution).

Reporting “unknown bacteria” is likely far more useful to a clinician than a “negative” result that would be returned from panels designed by specific sensors. This ability mirrors that of mass spectrometry and other fingerprinting systems, but the statistical underpinning could lead to theoretically grounded approaches with more research. Based on limitations in hard thresholding, our assay in its current form would more likely only be able to report “faulty solution” since “unknown bacteria” is a more specific call with a different clinical decision pathway.

## 3 Discussion

We demonstrate a new scalable framework for infection diagnostics that leverages the sparsity of samples and the Poisson distribution of microfluidic capture. Our nonspecific probes coupled with this framework enabled the quantification of 9 bacterial genera with 5 probes and 2 primers at levels as low as approximately 200 total copies of the 16S gene. The limit of quantification appears to be sufficient for many clinical applications, although our samples with only pre-extracted bacterial genomic DNA are highly idealized. Real-world samples with background nucleic acids are likely to hinder performance, but improvements in cluster separation through assay optimization may offset these effects.

More generally, we present theoretical results that substantially broaden the types of sensors that allow **λ** to be identifiable. However, our SPoRe algorithm is modular for *any* user-defined sensing function, and our theoretical conditions are sufficient but not necessary. We encourage users to proceed with simulations even if their sensing model is outside the scope of our currently developed theory. As a nonconvex optimization process, our approach is not guaranteed to find the best solution. New theoretical results that provide convergence guarantees and worst-case error quantification would substantially increase user confidence in this approach. For the time being, we have demonstrated a method for flagging solutions that are inconsistent with the observed data. This technique can help identify samples that contain analytes outside of the designed panel and solutions that have converged to poor local optima (if any). However, both the reliability of such flagging and the accuracy of estimated analyte quantities may be jeopardized if the model provided to SPoRe does not closely match real-world conditions.

The application of MLE to ddPCR data is not new, but our generalization offers unique insights and practical advantages for microfluidics-based diagnostics. These advantages stem from our tolerance of ambiguity from individual partition measurements. Conventionally, to achieve scalable diagnosis of many analytes, single-analyte capture is enforced by a limiting dilution (∑_*n*_ *λ*_*n*_ ⪅ 0.1) such that individual partition measurements can be directly classified. In diagnostics, sample concentrations are unknown and can vary over multiple orders of magnitude. We show how our framework tolerates multi-analyte capture such that samples of higher concentrations do not flood our system. The ability to diagnose with higher **λ** also reduces the need to generate many partitions. Enabling diagnostics with, for instance, hundreds of microchambers could enable scalable multiplexing on static, point-of-care formats. Asynchronous fingerprinting offers a second practical advantage. Many biosensing techniques are limited in *M*, the number of orthogonal measurements acquired from each partition, either due to spectral overlap or sensor cross-reactivity. However, more sensors may be necessary to assign unique fingerprints to each analyte. Asynchronous fingerprinting bypasses this limitation by enabling reconstruction of the signal from partial fingerprint measurements. Still, our previous theoretical and simulated results indicate that users should first maximize *M* and increase *G* as necessary [30].

Our ddPCR assay has a few critical limitations that warrant future development. First and foremost, we designed probes to assign unique barcodes to the bacterial genera in our chosen panel. However, future LNA probe design must account for bacteria outside the panel that could plausibly appear in a sample to ensure that the designed barcodes are specific to the intended pathogens. These design requirements motivate new bioinformatics tools. In general, we recommend that our framework is applied in applications where the scope of plausible analytes in the intended samples is well understood such that the specificity of the analytes’ fingerprints can be verified. Second, we amplified the full 16S gene for maximum sequence flexibility in probe design. Amplifying the full length, 1500 base pair gene is well beyond the manufacturer’s recommendations. In ddPCR, longer extension times and high cycle counts can generally only help improve endpoint amplitudes and clustering. Here, our PCR protocol alone was over 8 hours which is not ideal for infection diagnostics. Future iterations on our approach could either employ custom master mixes with faster polymerases, restrict the amplicon to a shorter 16S segment, or replace hard thresholding with probabilistic noise models at faster cycling conditions. Third, many clinical infections may be caused by bacteria or fungi. Multiplexing primers to include eukaryotic marker genes along with the 16S primers for bacteria could enable broadening the panel. All of the above lines of research should also consider the optimal marker genes to target with nonspecific probes; certainly, no single gene target will be optimal for all applications in microbial diagnostics.

While there is room for improvement in the ddPCR approach, expanding the measurement space with non-binary measurements would dramatically improve diagnostic performance as fewer sensors could assign unique fingerprints to microbes at a higher rate. Some of these non-binary effects may be readily present in the droplet “lean” and “lift” likely caused by weak interactions between probes and slightly mismatched amplicons. These potential sources of additional information were ignored by the hard thresholding employed in this work. Moreover, while we present our framework in the context of microbial diagnostics where our research group is most familiar, we note the generality of the approach for other diagnostic applications. Combining conventional sensors with new techniques in microfluidics and signal processing will offer a suite of new interdisciplinary approaches to scalable, multiplexed biosensing.

## 4 Methods

### 4.1 Theory: Identifiability with Common Types of Sensors

For estimating **λ**, the property of *identifiability* means that there is a one-to-one correspondence between each realizable distribution of measurements and the Poisson rates **λ**: if *p*(**y**| **λ**) = *p*(**y** |**λ**′) for all **y**, then **λ** = **λ**′. From an optimization perspective, identifiability implies that the **λ**^*^ is the unique global optimum to the likelihood function if we have infinite measurements. Therefore, identifiability is a necessary condition for our method to work.

#### 4.1.1 Notation

We use bold face upper and lower case letters for matrices and vectors, respectively. Non-bold, lower case letters represent scalars. We denote the vector of all zeros as **0** with its dimensionality dependent on context. We use script letters (A, B, etc.) to denote sets. We denote **e**_*j*_ as the standard basis vector with **e**_*j*_ = 1 and **e**_*i*_ = 0 for all *i* ≠ *j*. Let **a** and **b** be two arbitrary vectors of the same dimension, and let *a*_*i*_ and *b*_*i*_ denote their *i*th elements. We use supp(**a**) to denote the support of vector **a** defined as the index set where *a*_*i*_ > 0 for *i* ∈ supp(**a**). We use the notation **a** ⪰ **b** to imply that *a*_*i*_ ≥ *b*_*i*_ ∀*i*, and we use **a** ≻ **b** to further imply the existence of at least one index *i* where *a*_*i*_ > *b*_*i*_. A set in the subscript of a vector such as **x** _**𝒜**_ refers to the subvector of **x** indexed by the elements of 𝒜. We make frequent use of the shorthand **a** to denote the summation over elements of a vector **a**.

#### 4.1.2 Definitions and Assumptions

We treat the dataset of measurements from all partitions in a sensor group as samples of a random variable **y**. The signal (i.e., analyte quantities in a partition) **x** is *N* -dimensional with **x** ∼ Poisson(**λ**^*^). Each signal is measured by *M* sensors to yield the observation vector **y** (e.g., *M* fluorescence measurements). We define the function 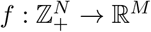 that is composed of *M* scalar functions 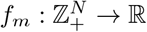. A particular measurement value *y*_*m*_ is determined by the sensor output *f*_*m*_(**x**) plus some additive, zero-mean random noise *n*_*m*_.

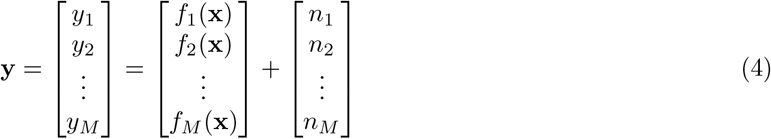

Note that the sensor functions *f*_*m*_ are group-dependent. For example, each group may have different probes.

We assume that our sensors are *monotonic* and that our analytes obey *responsiveness* and *fingerprint equivalence*. Given these properties, we prove sufficient conditions for identifiability. Without loss of generality, we will say that all *M* sensor functions are monotonic.

##### Definition 4.1

(Monotonic Sensors). *A sensor function f*_*m*_ : 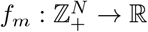 *is monotonic increasing if* **a** ⪰ **b** ⇒ *f*_*m*_(**a**) ≥ *f*_*m*_(**b**) *and monotonic decreasing if* **a** ⪰ **b** ⇒ *f*_*m*_(**a**) ≤ *f*_*m*_(**b**).

Monotonic functions are very common and natural; for instance, many sensing modalities have a monotonic increasing sigmoidal response to their input. Any time a Lemma or Theorem relies on monotonicity, its proof will assume all *M* sensors are monotonic increasing without loss of generality. Next, we define the *responsiveness* property of analytes:

##### Definition 4.2

(Responsiveness). *If f* (**e**_*i*_) ≠ *f* (**0**), *the analyte indexed by i is said to be responsive. If f* (**e**_*i*_) = *f* (**0**), *then the analyte indexed by i is nonresponsive. If we let* B *define the set of indices for all such nonresponsive analytes (i* ∈ B*), then for any two signals* **x** *and* **x**′, *f* (**x**) = *f* (**x**′) *if* 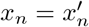 *for all n ∉* B.

In other words, a nonresponsive analyte does not influence the sensor output regardless of its quantity. An analyte is considered “responsive” if a single copy yields a different measurement than the null signal (e.g., an empty microfluidic partition).

We define a final intuitive condition on our system called *fingerprint equivalence*. The *fingerprint* of analyte *n* is the measurement yielded by an isolated copy of the analyte, or *f* (**e**_*n*_). Among analytes with identical fingerprint responses within a sensor group, the total number of occurrences of these analytes dictates the output response. In other words, the sensors treat these analytes as interchangeable copies of each other.

##### Definition 4.3

(Fingerprint Equivalence). *Let 𝒳* ⊆ {1, …, *N* } *be an index set of analytes with identical fingerprints, i*.*e. f* (**e**_*i*_) *is fixed for all i* ∈ 𝒳. *A system h as the fin gerprint equivalence property if for any pair of vectors* **x** *and* **x**′ *with* supp(**x**), supp(**x**′) ⊆ X *and* 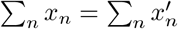, *we have f* (**x**) = *f* (**x**′).

Note that even if all analyte fingerprints are distinct, multiple signals can still map to the same measurement vector since we allow for cases of multi-analyte capture in the same partition. We define these signals as members of a *collision set*.

##### Definition 4.4

(Collision Sets). *The collision set 𝒞*_**x**_ *for signal* **x** *is the set of all signals* **x**′ *that satisfy f* (**x**′) = *f* (**x**).

We define 𝒰 as the set of unique collision sets. In 2-channel ddPCR with binarized measurements, there are four collision sets in each sensor group 𝒰 (0, 1 ^2^). It will soon be clear that observations **y** are drawn from a mixture model. We can define each *mixture element* as follows:

##### Definition 4.5

(Mixture Element). *The mixture element ℰ*_**x**_ *for signal* **x** *is the set of all signals* **x** ′ *that satisfy p*(**y**|**x**) ∼ *p*(**y**|**x**′).

Note with any zero-mean noise, *p*(**y**|**x**) ∼ *p*(**y**|**x**′) ⇒ *f* (**x**) = *f* (**x**′) such that ℰ_**x**_ ⊆ 𝒞_**x**_. In some cases, such as additive white Gaussian noise, ℰ_**x**_ = 𝒞_**x**_. We define 𝒱 as the set of unique mixture elements with arbitrary ℰ_*v*_ ∈ 𝒱.

#### 4.1.3 Proof of Identifiability

With *G* different sensor groups indexed by *g*, we assume that the sensor group applied to a measurement **y** is known and deterministic. Each sensor group has a different function *f* that maps **x** to *M* -dimensional space (e.g., different probes in ddPCR). Identifiability means that *p*(**y**| **λ**) = *p*(**y** |**λ**′)∀**y**, *g* ⇒ **λ** = **λ**′. Each **λ** must yield a unique set of *G* distributions of measurements.

We will use the notation 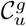 and 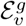 to specify the group *g* when necessary. For an arbitrary group, we can express *p*(**y**|**λ**) as:

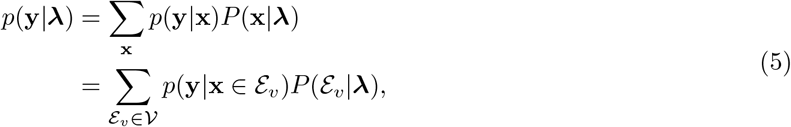

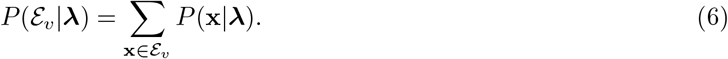

If a mixture distribution is identifiable, it means that identical distributions must come from the same set of weights on the mixture elements; in this context, *p*(**y** |**λ**) ∼ *p*(**y** |**λ**′) *P* (ℰ_*v*_ |**λ**) = *P* (ℰ_*v*_ |**λ** ′)∀*v*. Many finite mixtures (what we practically have in MMVP) and countably infinite mixtures with common noise distributions are identifiable [40, 41], and we assume that the system noise characteristics lend to an identifiable mixture. However, we need to prove the identifiability of MMVP, or that equal mixture weights implies equal Poisson parameters: 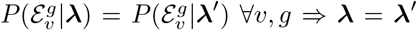. Note that because 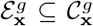 and unique collision sets are disjoint, 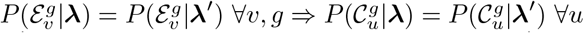.

We assume 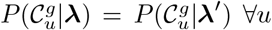 and prove the implication of **λ** = **λ**′ given a set of monotonic sensors and with analytes exhibiting responsiveness and fingerprint equivalence in all *G* groups. We first focus on what can be concluded from a single, arbitrary sensor group (dropping the *g* superscript) with analytes potentially having nonunique fingerprints, and then we conclude with how multiple groups can be pooled to achieve identifiability. Again, note that our analysis will focus on monotonic increasing sensors without loss of generality.

Define A ⊆ {1, …, *N* } such that analytes indexed by *a* ∈ 𝒜 are all responsive such that there exists some *m* such that *f*_*m*_(**e**_*a*_) > *f*_*m*_(**0**). Define the complementary set ℬ with nonresponding analytes indexed by *b*.

##### Lemma 4.1.

*If f*_*m*_ *is monotonic increasing for all m* ∈ {1, …, *M* }, *and if only the analytes indexed by a* ∈ 𝒜 ⊆ {1, …, *N* } *are responsive, then f* (**x**) = *f* (**0**) *if and only if* **x**_**𝒜**_ = **0**.

*Proof*. Consider **z** such that **z**_**𝒜**_ = **0**. Note that supp(**z**) ⊆ B. Because analytes indexed by *b* ∈ ℬ are nonresponding, *f* (**z**) = *f* (**0**) by definition. Next, we prove the forward condition, *f* (**x**) = *f* (**0**) ⇒ **x**_**𝒜**_ = **0**, by contradiction. Say *f* (**z**) = *f* (**0**) and let **z** satisfy *z*_*a*_ ≥ 1 for some *a* ∈ 𝒜. By definition of 𝒜, *f* (**e**_*a*_) > *f* (**0**), and **z** ⪰ **e**_*a*_. With monotonic functions, *f* (**z**) ⪰ *f* (**e**_*a*_) ≻ *f* (**0**) and we have arrived at a contradiction.

The key concept to carry forward is that values of elements in **x**_*B*_ are entirely arbitrary for the analysis of collision sets.

##### Lemma 4.2.

*Let f*_*m*_ *be monotonic increasing for all m* ∈ {1, …, *M* }, *a nd let only the analytes indexed by a* ∈ 𝒜 ⊆ {1, …, *N* } *be responsive. If P* (𝒞_**0**_|**λ**) = *P* (𝒞_**0**_|**λ**′), *then* 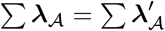

*Proof*. By Lemma 4.1, C_**0**_ contains all **x** with **x** _**𝒜**_ = **0** with arbitrary values on **x**_*B*_. Therefore, *P* (C_**0**_|*λ*) = *P* (C_**0**_|*λ*′) implies

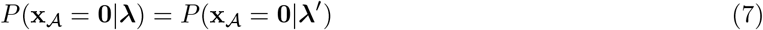

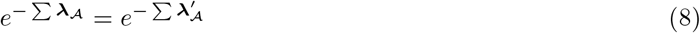

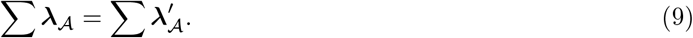

##### Lemma 4.3.

*Let f*_*m*_ *be monotonic increasing for all m* ∈ {1, …, *M* }, *and let only the analytes indexed by a* ∈ 𝒜 ⊆ {1, …, *N* } *be responsive. For a* ∈ 𝒜, *if for all* 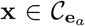 *is the only nonzero value in* **x**_*A*_, *then* 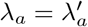

*Proof*. We assume 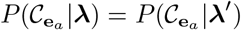. By definition of A, *f* (**e**_*a*_) ≻ *f* (**0**). If *x*_*a*_ is the only nonzero value of *x* _*𝒜*_, we have 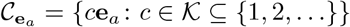. Then, 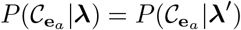 implies

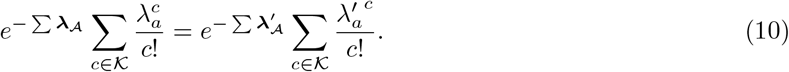

Using Lemma 4.2, in 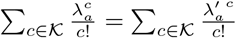, which implies 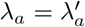 since the function on both sides is monotonic

From here, we first derive results for the special case where all analytes indexed by 𝒜 have unique single-copy fingerprints. Afterwards, we generalize to multiple groups, allowing for equal nonzero fingerprints within a group. The next Lemma guarantees at least one index *a* to which Lemma 4.3 can be applied.

##### Lemma 4.4.

*Let f*_*m*_ *be monotonic increasing for all m* ∈ {1, …, *M* }, *and let only the analytes indexed by a* ∈ 𝒜 ⊆ {1, …, *N* } *be responsive. If f* (**e**_*i*_) ≠ *f* (**e**_*j*_) ∀*i, j* ∈ 𝒜 *with i* ≠ *j*, ∃*a* ∈ 𝒜 *such that all* 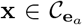 *are nonzero in* **x** _**𝒜**_ *only on index a*.

*Proof*. First, with unique nonzero fingerprints in A and monotonic sensors, the fingerprint responses *f*_*m*_(**e**_*a*_) can be sorted. Starting arbitrarily with *m* = 1, we can select the minimal set ℳ ⊆ A that minimizes *f*_1_(**e**_*a*_) such that ∀*a* ∈ ℳ, ∀*j* ∈ ℳ_*c*_, *f*_1_(**e**_*a*_) *< f*_1_(**e**_*j*_). If | ℳ | > 1, then the process can be repeated with *m* = 2 (and so forth) on the subset ℳ until there is one unique minimum and its corresponding index *a*.

For this **e**_*a*_, all *i* ∈ 𝒜 \ {*a*} satisfy *f*_*m*_(**e**_*i*_) > *f*_*m*_(**e**_*a*_) for at least one *m*. Therefore, signals in the collision set 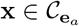 must satisfy **x**_*i*_ = 0 for all *i* ∈ 𝒜 \ {*a*}. Signals with at least one **x**_*i*_ ≥ 1 would have at least one *m* where *f*_*m*_(**x**) > *f*_*m*_(**e**_*a*_), and therefore not be in the collision set by definition. This conclusion completes the proof by contradiction.

Next, we show how this result chains to all analytes indexed in 𝒜.

##### Lemma 4.5.

*Let f*_*m*_ *be monotonic increasing for all m* ∈ {1, …, *M* }, *and let only the analytes indexed by a* ∈ 𝒜 ⊆ {1, …, *N* } *be responsive. If f* (**e**_*i*_) ≠ *f* (**e**_*j*_) ∀*i, j* ∈ 𝒜 *with i* ≠*j*, 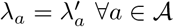.

*Proof*. Lemmas 4.3 and 4.4 guarantee at least one *a* that yields 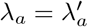. Let us call this index *a*_1_ and define the subset 𝒮 ⊆ 𝒮, the subset of indices for which 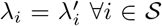. At this point, 𝒮 = {*a*_1_}. Repeating the process in the proof of Lemma 4.4, we can find a new index *a*_2_ that satisfies 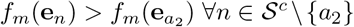 for at least one *m*.

For the direct proof of identifiability, we assume 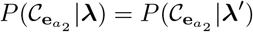, or:

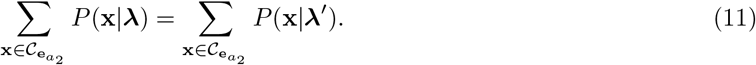

Among signals **x** in 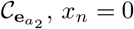 for *n* ∈ 𝒮 \ {*a*_2_} because 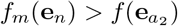 for some *m*, and sensors are monotonic. These signals can also be partitioned into those with 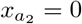, and those with 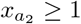.If any of the former type exist, then *x*_*i*_ > 0 for some of the indices *i* ∈ 𝒮. For instance, we could have 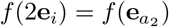 with *x*_*i*_ = 2. These signals’ terms in the summation follow the form 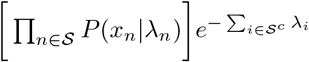. Note that 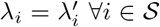 combined with Lemma 4.2 yields 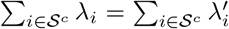. Because 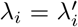 for *i* ∈ 𝒮, the product component is equal on both sides as well. Therefore, these terms can be eliminated in Equation (11). We will denote the set of remaining **x** in the summation as 𝒞*′*.

In 𝒞*′*, we have 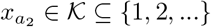. Therefore:

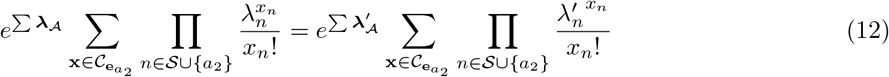

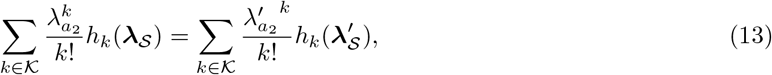

where *h*_k_ is the mapping 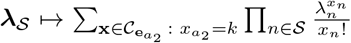. Because 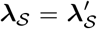, we can replace both *h*_*k*_ (**λ** _*𝒮*_) and 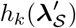 by constants *H*_*k*_. Therefore,

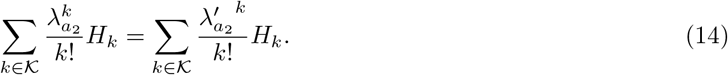

Because *H*_*k*_ ≥ 0, both sides are monotonic in 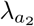 such that 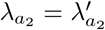. Now, *a*_2_ can be added to 𝒮 and the process can be repeated until 𝒮 = 𝒜 such that 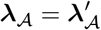.

Now, we will extend this result to the case of having equal fingerprints in the same sensor group—i.e., that *f* (**e**_*i*_) = *f* (**e**_*j*_) for some pairs *i, j*.

##### Lemma 4.6.

*Let f*_*m*_ *be monotonic increasing for all m* ∈ {1, …, *M* }, *and let only the analytes indexed by a* ∈ 𝒜 ⊆ {1, …, *N* } *be responsive. Define the disjoint sets 𝒜*_1_, *𝒜*_2_, … *𝒜*_*C*_ *indexed by c with* 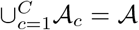 *such that for all i, j* ∈ 𝒜_*c*_, *f* (**e**_*i*_) = *f* (**e**_*j*_). *Then*, 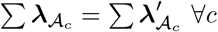.

*Proof*. Note that the Poisson distribution has the property that if *x*_*i*_ are each independently drawn from Poisson(*λ*_*i*_), then ∑ *i* ∈ 𝒜 _*c*_ *x*_*i*_ ∼ Poisson 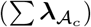. We can then simply define a dummy variables 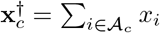 and 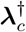 such that 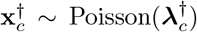. This dummy variable represents an “analyte” that appears with a distribution governed by the total quantities of analytes with the same fingerprint. However, what matters for identifiability is the sensor functional values of signals, i.e. that *f* (**a**) = *f* (**b**) if for all 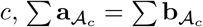. Namely, 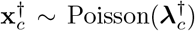 by fundamental properties of the Poisson distribution, but it is only with the condition of fingerprint equivalence (Definition 4.3) that lets us apply all previous results that are based on collision sets, i.e., sets of signals with equal functional values. These yield 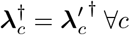, or 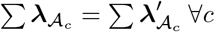.

##### Theorem 4.7.

*Let g index G different sensor groups that satisfy fingerprint equivalence and that contain monotonic, saturating sensors. For each group g, define the row vector* 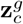 *of zeros and ones with ones in the indices associated with 𝒜*_*c*_. *Define the N -column matrix* **Z**_*g*_ *whose C rows are comprised of* 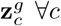. *Define the N -column matrix* **Z** *as the vertical concatenation* **Z**_*g*_ ∀*g. If* rank(**Z**) = *N, then* **λ** = **λ**′.

*Proof*. This theorem is a formal way of saying that Lemma 4.6 must yield *N* independent equations when applied to all groups where the sensing and system conditions hold. We can consider the system of equations yielded by Lemma 4.6 and represented by **Z*λ*** = **Z*λ***′, or **Z**(**λ** − **λ**′) = **0**. If rank(**Z**) = *N*, then it follows that **λ** = **λ**′. Therefore, we have 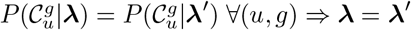, concluding our proof of identifiability.

### 4.2 Bacterial Panel

We ordered bacterial species’ genomic DNA from the American Type Culture Collection (ATCC, Manassas, VA). The species’ names and their ATCC identifiers are as follows: *A. baumannii* (BAA-1605), *B. fragilis* (25285), *E. cloacae* (13047), *E. faecium* (BAA-23200, *E. coli* (11775), *K. pneumoniae* (13883), *P. aeruginosa* (BAA-1744), *S. aureus* (12600), *S. epidermidis* (14990), *S. saprophyticus* (15305), *S. agalactiae* (13813), and *S. pneumoniae* (33400). Particular strains were selected based on their availability at the time of purchase and only if ATCC provided whole genome sequence information for the isolate. DNA was resuspended and aliquoted according to ATCC’s instructions at approximately 10^6^ genome copies per microliter. DNA aliquots were stored at -4 C until use.

### 4.3 Probe Design

All oligonucleotides were acquired from Integrated DNA Technologies (Coralville, IA) with HPLC purification and are listed in Table 1. We used ThermoBLAST from DNA Software (Plymouth, MI) to align the 16S primers (27F and 1492R from [11]) against bacterial genomes and find amplicons. We passed these amplicons to a custom Matlab script to design probes. We needed probes with a melting temperature (*T*_*m*_) a few degrees higher than that of the primers, targeting at least 65°C. Probes for barcoding must hit multiple bacterial taxa, and shorter probes are naturally less specific. LNA’s increase probe melting temperature in a highly position-specific manner. To avoid combinatorially increasing our probe search space, we deferred LNA positioning until after sequence selection. We used heuristics to filter for probes that would have sufficient *T*_*m*_ with some flexibility in LNA positioning. We chose a sequence length of 11 nucleotides with 5-8 GC nucleotides, without four consecutive G’s or C’s, and without a G on the 5’ end to avoid self-quenching of the fluorophore. We used Smith-Waterman alignment in Matlab to pre-screen for probes that self-hybridize and to assess cross-hybridization of probes amongst the evolving candidate set. To assist in achieving near binary measurements, we considered perfect matches on all 16S copies to be “1” for a genome, and for imperfect homology, we filtered for sequences that had neither nine consecutive matches nor a single G-T mismatch. This latter filtering is a proxy for ensuring that probes have weak, negligible interactions against 16S sequences where they do not have perfect complementarity. The former filtering for positive hits was intended to avoid the issue of mixtures of barcodes for any particular bacteria for simplicity in our initial demonstration.

**Table 1:**
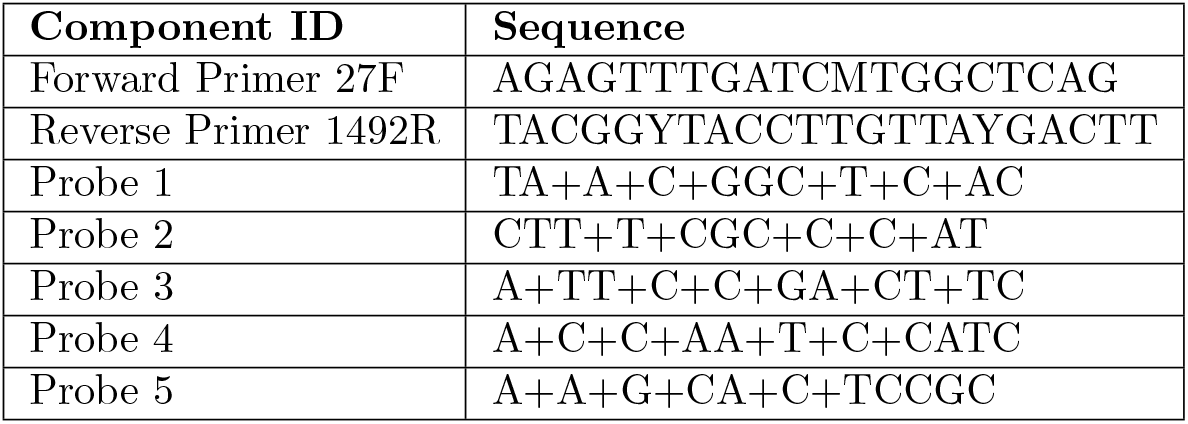
Oligonucleotides used in this study. A “+” preceding a base indicates that the LNA analogue of the base was used. Probes 1, 2, and 5 were tagged with HEX and Probes 3 and 4 with FAM on the 5’ ends. The dark quencher Iowa Black FQ was placed at the 3’ ends.

Given a set of filtered, candidate probes, we used a coordinate ascent strategy to iteratively optimize a set. We hypothesized that barcoding the full-length 16S gene with probes could achieve genus level resolution, as sequencing the full gene achieves a mix of genus and species resolution. As a result, we encouraged similarity of the three *Staphylococcus* species and the two *Streptococcous* species. Define 𝒮 as a set of pairs of bacteria (*b*_*i*_, *b*_*j*_) within a genus that are similar. The complementary set 𝒟 includes all other bacterial pairs. Let **k**_*p,I*_ represent the 11-mer barcode of bacteria *i* with probes indexed by *p*. Coordinate ascent sought to solve:

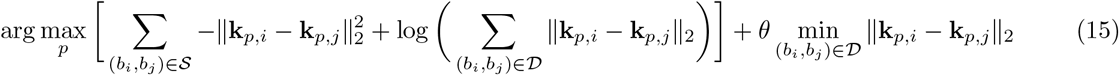

The first term is taken from research in metric learning [42], and the second term (with a weight of *θ* = 10) highly rewards some nonzero separation between all bacterial pairs that are intended to be discriminated between. We chose an initial random set of probes that passed our cross-hybridization check. We iteratively cycled through a shuffled order of the candidate set of probes, evaluating one probe at a time for replacement with any of the other probes that passed the initial filtering step. If replacing a probe improved the objective function, the probe set was updated and the search continued. The algorithm terminated when all probe sequences had been evaluated for replacement but not replaced. For the chosen set of sequences, we evaluated the alignment against 16S genes with imperfect homology (the zeroes in the barcodes). As much as possible, we positioned LNAs at mismatch sites to improve the thermodynamic discrimination against these sequences. We evaluated all *T*_*m*_’s in IDT’s OligoAnalyzer, positioning additional LNAs as necessary to reach a sufficient probe *T*_*m*_. Final probe sequences are listed in Table 1.

### 4.4 Preparation of Bacterial Samples

First, we prepared monomicrobial dilutions of genomic DNA in MilliQ purified water. One dilution was prepared for each bacteria, approximately targeting a concentration **λ** between 0.2 and 2 (“Concentration 1”). We diluted each of these by 1/2 to yield a second dilution of “Concentration 2.” We used a custom script to assign random combinations of these bacterial dilutions to samples, generating five samples with *k* = 2 unique bacteria and six samples of *k* = 3 and *k* = 4. We reserved one sample to be a no template control (NTC, water alone). The probability of drawing each bacteria was adaptively weighted to encourage approximately even representation of each taxa across the samples (note, *Staphylococcus* and *Streptococcus* species were lumped and treated as one taxa each to not overrepresent their constituent species).

### 4.5 Droplet Digital PCR

Primers were at 900 nM and all probes were at 125 nM. We used Bio-Rad’s ddPCR Multiplex Supermix and prepared master mixes, generated droplets, and read out droplets according to the manufacturer’s instructions. For PCR cycling, extension times were set to 7 minutes because of the long amplicon (approximately 1500 base pairs) that is atypical in ddPCR, partly following guidance from [43] and internal data (not shown). PCR cycling was as follows: 95°C for 10 minutes (initial denaturation and hot-start deactivation), 60 cycles of 94°C for 30 seconds (denaturation) and 60°C for 7 minutes (anealing and extension), 98°C for 10 minutes. Ramp rates during cycling were set to 2°C/second. Samples were refrigerated at 4°C for 30 minutes prior to droplet readout.

### 4.6 Plate Arrangement and Manual Thresholding

Figure 9 illustrates how we arranged the samples in our ddPCR plate. The first two rows serve as reference samples, allowing us to estimate the ground truth concentration of the two dilutions based on the proportion of empty droplets. Empty droplets were determined by manual thresholding. These are also color coded to reflect which group mixture was applied to each. The amplitudes of positive droplets in the reference rows, and their anticipated partial barcodes, were used to guide manual thresholding cutoffs for the rest of the plate. Each polymicrobial sample was distributed across four wells each assigned to a probe group (“G1” for Group 1, etc.).

**Figure 9:**
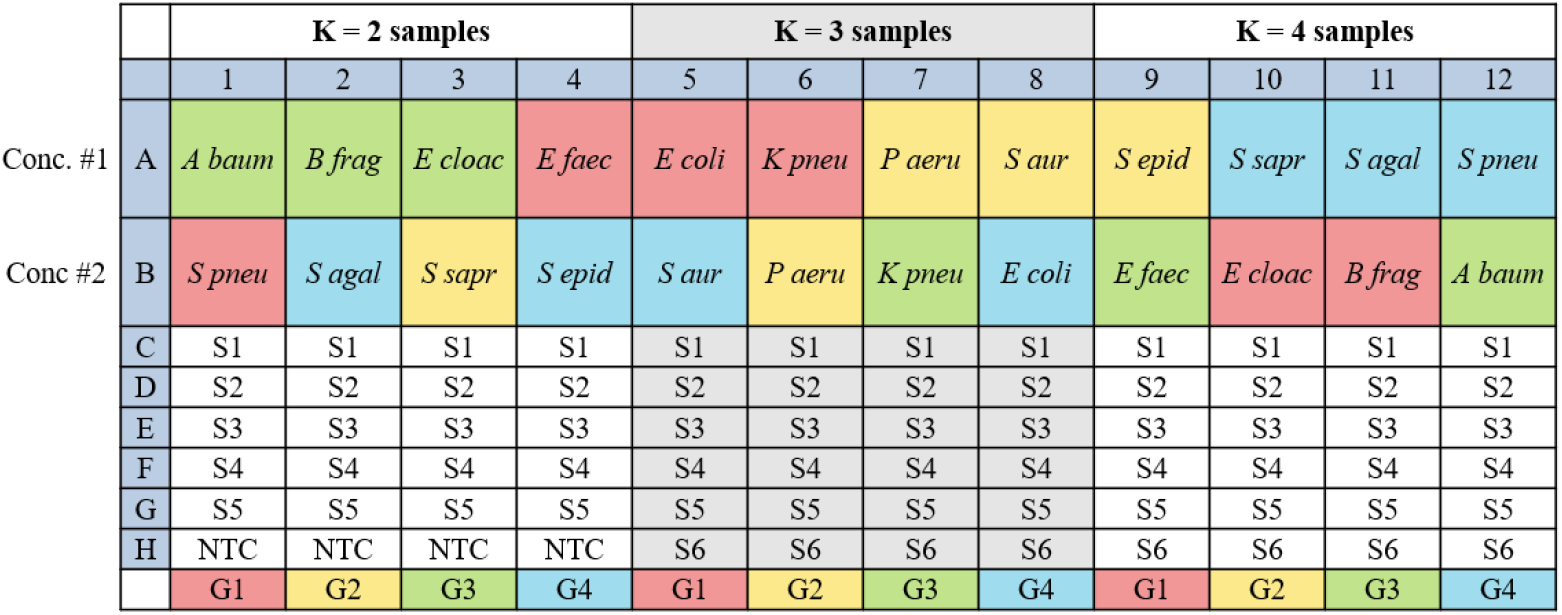
Plate layout for ddPCR test samples. The first two rows served as references for ground truth concentration estimation of monomicrobial dilutions and manual thresholding of all wells. The colors of the wells in rows A and B correspond to the probe group applied. Random mixtures of bacteria were distribtued across the rest of the plate with each mixture being applied to four wells, each with a different subset of two probes defining the 16S barcodes.

**Figure 10:**
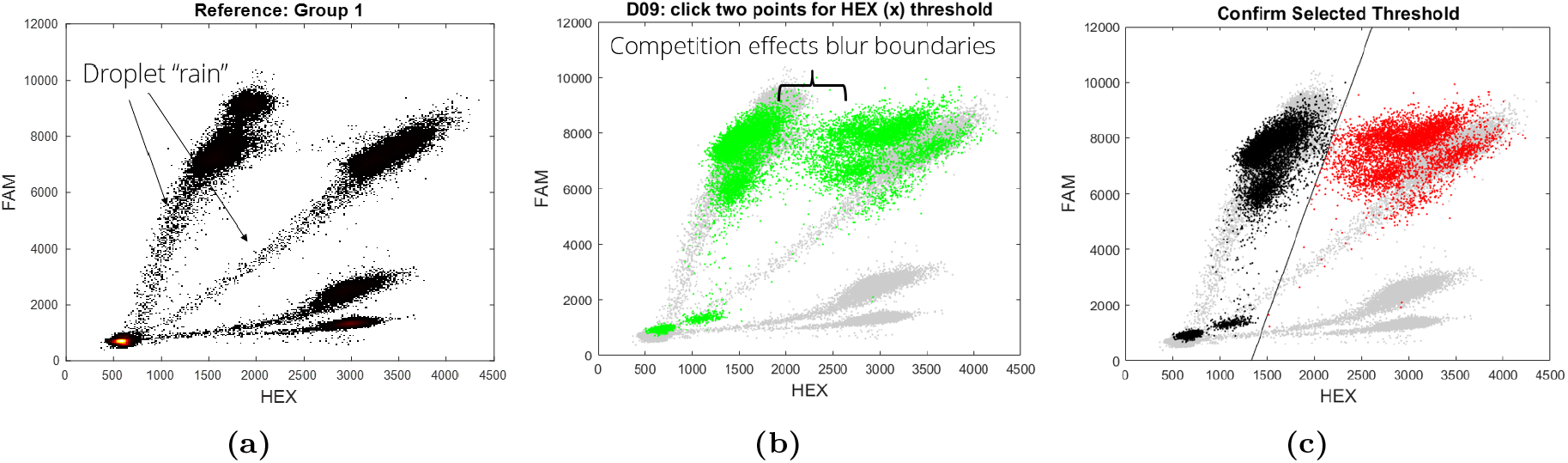
Example process for manual thresholding with noted challenges. **(a)** All reference data coming from the same probe group was pooled and displayed to serve as a visual reference. Droplet “rain” is evident in each cluster. **(b)** Raw data from a polymicrobial sample was overlaid on the reference data with the same corresponding probe group. An example is shown with the raw data from D09 (Group 1, *k* = 4 sample number 2) overlaid with the Group 1 reference data. PSC effects, as expected, create subpopulations of droplet measurements between the binary clusters. We speculate that the small additional cluster near zero is due to droplets where probes partially interacted with amplicons due to imperfect sequence homology. **(c)** After the user selects two points to define a line for thresholding, the plot is updated to allow the user to visually confirm the results. Red points are assigned the value 1, and black points are assigned 0.

For manual thresholding, we pooled the data from the reference wells for each group and overlaid a particular well’s data (Fig. 10a,b). Due to some mild “lean” and “lift” of the raw ddPCR clusters caused likely by partial probe interactions, we allowed any linear threshold for each fluorescence channel determined by two user-selected points. The user confirmed the threshold selection (Fig. 10c). This process was repeated for all wells, including the reference wells, and conducted twice for each well to define thresholds for both HEX and FAM. Ultimately, all droplets from each well are converted to two dimensional binary measurements.

### 4.7 Settings for SPoRe

We set the initial value of **λ** to the constant vector **0.1** for an unbiased initialization. We performed gradient descent to minimize the negative log likelihood with backtracking line search to adaptively tune the learning rate at each iteration. First, the gradient is computed and applied to the estimate **λ** with an initial learning rate, and *λ*_*n*_ is projected to max(*λ*_*n*_, 10^−6^). Values of *λ*_*n*_ = 0 cause numerical issues in gradient computations, hence the small offset. SPoRe checks if the negative log likelihood decreased given the new **λ**. If it did not, the iteration’s step size was cut in half and the process is repeated. SPoRe only commits to an update to **λ** after confirming a reduction in the negative log likelihood, at which point it resets the learning rate and proceeds to the next iteration. SPoRe terminates when the log likelihood’s relative change over the previous five iterations falls below 10^−6^. In contrast to our original implementation of SPoRe [30], we use an exact gradient computation over the entire dataset made feasible by the nature of ddPCR data (Section 5.2) rather than a Monte Carlo approximation over a batch of data.

### 4.8 Assessing *E. cloacae* Barcode Variability

*E. cloacae*’s amplicons appeared to always interact with Probes 3 and 4, but a small subcluster appeared to lack the HEX response to Probe 1 (Fig. 11a). We used our SPoRe algorithm to estimate the barcode abundances in reference wells A3 and B10 (Fig. 9) which both contained Probe 1. After manual thresholding, the binarized data and “analytes” with barcodes [0, 1], [1, 0], and [1, 1] were passed to SPoRe, and SPoRe estimated the abundances of the amplicons with these responses. In this case, the [1, 0] quantity was nearly zero, consistent with the expectation that *E. cloacae* always interacted with a FAM probe. The fraction of amplicons with the HEX probe was determined by *λ*_[1,1]_*/*(*λ*_[1,1]_ + *λ*_[0,1]_). In A3 and B10, these were estimated to be 0.832 and 0.828, respectively. Our sequence analysis found eight 16S copies in the *E. cloacae* genome, so it is possible that one amplicon had a sequencing error such that Probe 1 truly binds to 7/8 copies. Therefore, column 3 of matrix **C** had 0.875 of barcode 3 and 0.125 of barcode 1 (Fig. 12).

**Figure 11:**
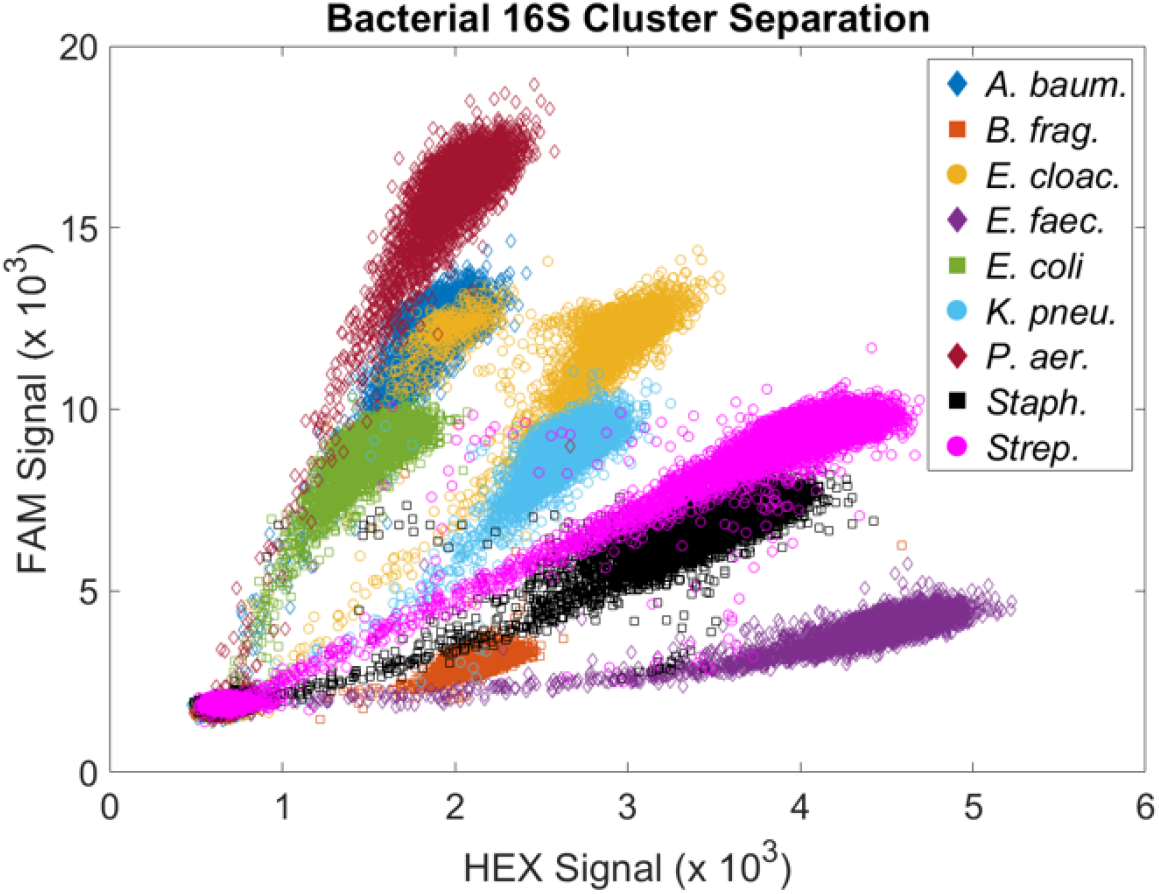
Separation of bacterial barcodes with amplitude multiplexing. Each cluster here is from a separate ddPCR reaction with one bacterial species in it. Data from the three *Staphylococcus* bacteria and the two *Streptococcus* bacteria were combined in this plot

**Figure 12:**
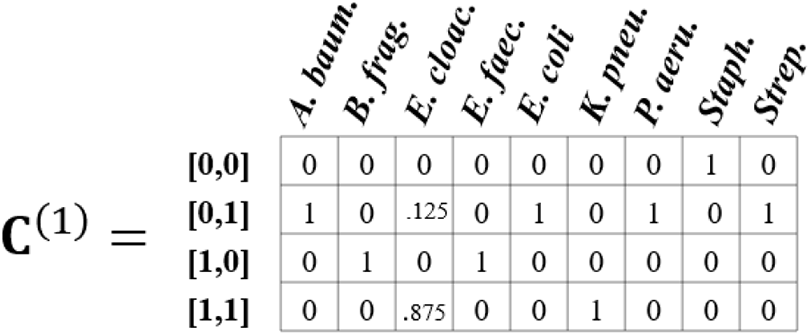
Example of partial barcode matrix for Group 1

**Figure 13:**
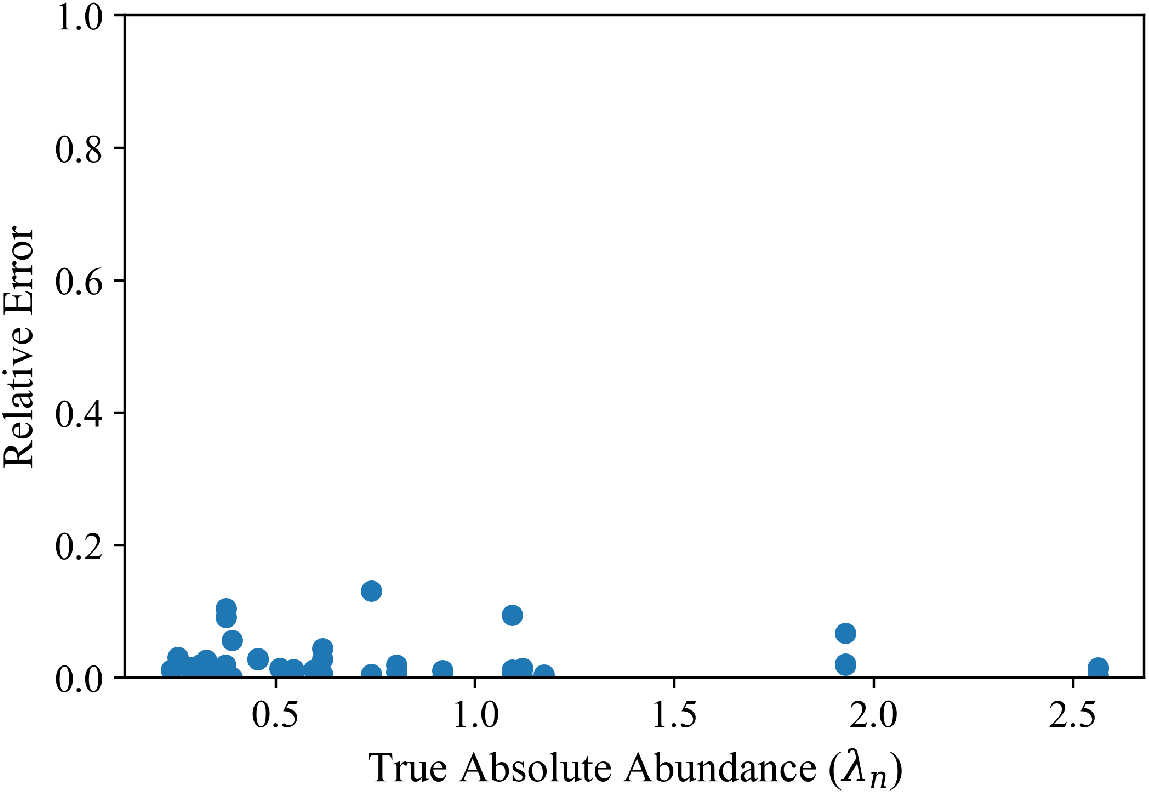
SPoRe’s performance on simulations of experimental concentrations. Given the estimated ground truth concentrations (**λ**^*^), we simulated binary measurement data to pass to SPoRe. SPoRe returns virtually perfect results with mean cosine similarity of 0.9997. Here, we plot the relative error in estimates of the absolute abundance for each bacteria, represented by 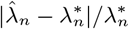 when running SPoRe on the simulated measurements.

## 5 Appendix

### 5.1 Amplitude Multiplexing with More than Two Probes

Amplitude multiplexing is a technique to resolve more probes than the available number of color channels, but it is typically used with each unique probe participating in an orthogonal assay with its own primer pair [35]. Probe concentrations can be adjusted to “move” the cluster positions. Here, we adjust probe concentrations with a single pair of primers. Probes 1 and 5 were tagged with HEX, and Probes 2-4 were tagged with FAM (Figure 3a). We tested a few concentration variations (results not shown), and the data depicted here is with Probes 1, 2, and 4 at 125 nM, and with Probes 3 and 5 at 250 nM. Based on each 16S gene’s barcode, droplets containing that gene will position in clusters whose channel intensities roughly correlate with the total probe concentration tagged with the corresponding fluorophore.

### 5.2 Exact Gradient Computation and *p*(y|x) Model for ddPCR

We first focus on the gradient resulting from a single probe group. In a single group, there are only four viable measurements with **y** ∈ **{**0, 1} ^2^. Let us define 𝒴= {0, 1} ^2^, and *p*_**y**_ as the proportion of the *D* measurements that equal **y**. We can then re-express Equation (2) as:

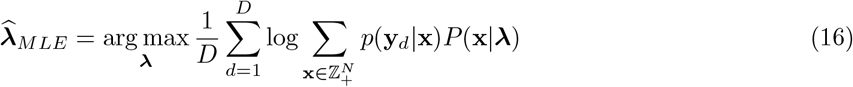

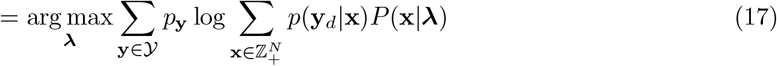

Here, we will use ***λ ≡ λ***^(*bact*)^ since we will be optimizing over the bacterial concentrations directly. In our case, *E. cloacae* is the only bacteria with a fractional abundance of a probe binding site - approximately 87.5% of its copies interact with probes 1, 3, and 4, and 12.5% interact with only probes 3 and 4 (Section 4.8). Similarly to how we defined **C** in Section 2.1, we can define **C**^(*g*)^ for group *g* with each bacterium’s fractional abundances of genes with a corresponding barcode. Figure 12 shows an example of **C**^(1)^, which can be generated with Figure 3a as a reference.

We define *p*(**y**|**x**) = ∏ _*m*_ *p*(*y*_*m*_|**x**). For *p*(*y*_*m*_ = 1|**x**), then *p*(*y*_*m*_|**x**) = 1 if the droplet has at least one copy of a gene that interacts with probe *m* and *p*(*y*_*m*_ |**x**) = 0 otherwise. For *p*(*y*_*m*_ = 0 |**x**), this likelihood is 1 if none of the genes in the droplet interact with probe *m* and 0 otherwise.

However, with the analyte current defined as a copy of the *n*th bacterium’s 16S gene, we must be careful. For instance, with index 3 corresponding with *E. cloacae*, if *x*_3_ = 1 in **x**, *p*(*y*_1_ |**x**) may not be 1 since one copy of *E. cloacae*’s 16S gene is not guaranteed to interact with probe 1. To resolve this, we will temporarily transform the problem to the space of gene barcodes for this group: 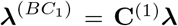. Note 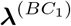 is 4-dimensional. We can define 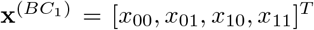 as the vector representing the quantities of 16S genes from any source bacteria that interact with the probes in the pattern noted in the subscript, noting 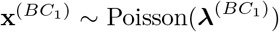. Lastly, let us define _**y**_ as that set where if 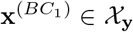, then 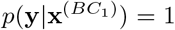.

Now we can rewrite Equation (17) as:

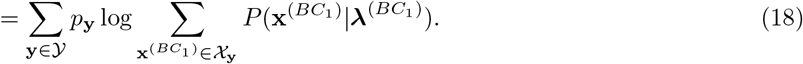

The linearity of gradients allow us to treat this one **y** at a time, summing the contributions from each **y** at the end. In general, treat 00 as short for [0, 0], 01 for [0, 1], etc. Let 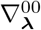 be the component of the gradient from **y** = [0, 0], 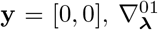 from **y** = [0, 1], etc. We will similarly define the mean log likelihood contributions as *ℓ*^00^, *ℓ*^01^, etc. Similarly, define the rows of **C**^(1)^ as 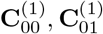, etc. By convention, **λ** and other vectors should be assumed to be column vectors, but the rows of **C**^(1)^ are row vectors. Thus we have:

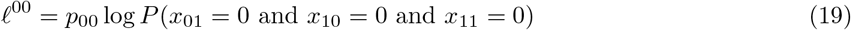

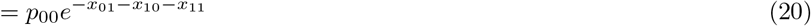

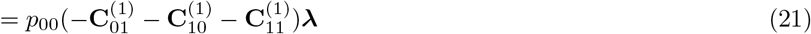

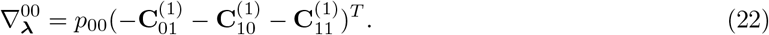

In the first line, we define the conditions for 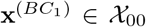 and solve. Genes that interact with either probe cannot be in droplets that yield **y** = [0, 0]. Next, for the **y** = [0, 1] response, at least one gene that interacts with the 2nd (FAM) probe must be present, and genes that interact with the HEX probe must be absent.

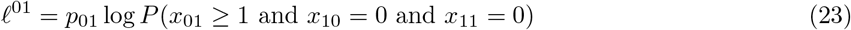

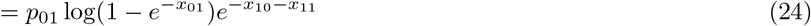

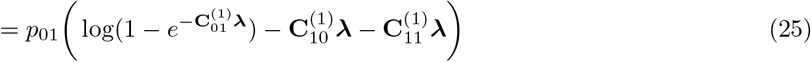

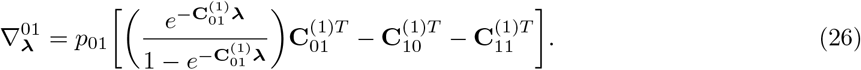

A virtually identical simplification for 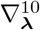 is omitted here. Lastly, for **y** = [1, 1], we have:

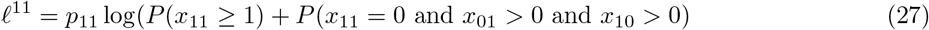

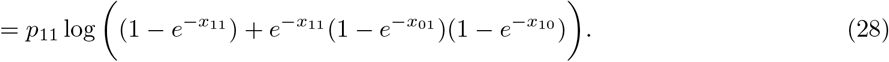

The remaining algebraic steps are omitted, but the final result is

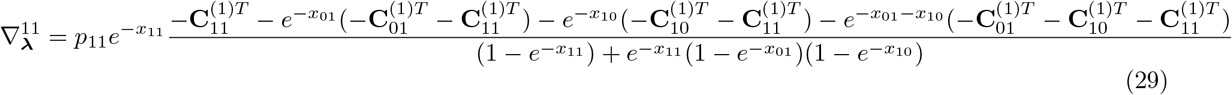

where 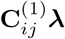 can be substituted for any *x*_*ij*_.

We can now say that for group 1, 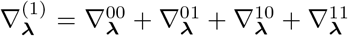. The above process can be repeated for any group *g*. Therefore, the final gradient vector (arbitrarily scaled) is

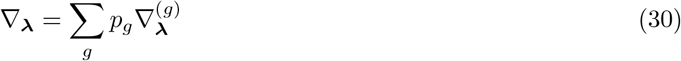

where *p*_*g*_ is the proportion of total droplets that come from group *g*.

### 5.3 Special Cases of MLE with ddPCR

From Equation (3) describing the generalized gradient in MLE, we consider two commonly employed special cases. First, if samples are sufficiently dilute such that partitions are either empty (**x**_*d*_ = **0**) or have only one analyte, the goal is often to identify each nonzero signal independently with a classification process. In other words, assays must be designed such that *p*(**y**_*d*_|**x**) > 0 *only* for 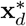 - the measurements are *unambiguous*. Setting the gradient equal to zero and simplifying leads to 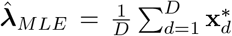. In practice, clusters of classes must have reliable decision boundaries and concentrations are estimated by totaling the classification results.

The second specialized case is common for ddPCR where for each PCR assay is specific for a target analyte and assigned to a particular channel. With *M* channels, *N* = *M* and each measurement unambiguously determines the presence or absence of each target sequence. Precisely, *p*(*y*_*d*_|**x**) is one or zero, and considerations of each analytes’ quantity *x*_*n*_ can be simplified to *x*_*n*_ = 0 (absent) or *x*_*n*_ > 0 (present). Each analyte *n* can be inferred independently such that 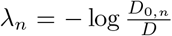 where *D*_0,*n*_ is the number of droplets that do *not* contain analyte *n*. This formula can be found by applying the above assumptions, setting Equation (3) to zero, and simplifying.

### 5.4 SPoRe’s Performance on Simulated Equivalents of Experimental Data

## Acknowledgments

This work was supported by NSF grant CBET 2017712 and the Rice Institute of Biosciences and Bioengineering. P.K.K. was supported by the NLM Training Program in Biomedical Informatics and Data Science (T15LM007093). D.L. and R.G.B. were supported by NSF grants CCF-1911094, IIS-1838177, and IIS-1730574; ONR grants N00014-18-12571, N00014-20-1-2534, and MURI N00014-20-1-2787; AFOSR grant

FA9550-22-1-0060; and a Vannevar Bush Faculty Fellowship, ONR grant N00014-18-1-2047.

## Notes

### Competing Interest Statement

Rice University has filed a patent application related to compressed sensing with microfluidic partitioning on which RD, RGB, PKK, DL, and HV are co-inventors. PKK has since founded a startup, Anvil Diagnostics Inc., which seeks to commercialize relevant technologies.

